# Differentiation of related events in hippocampus supports memory reinstatement in development

**DOI:** 10.1101/2023.05.25.541743

**Authors:** Nicole L. Varga, Hannah E. Roome, Robert J. Molitor, Lucia Martinez, Elizabeth M. Hipskind, Michael L. Mack, Alison R. Preston, Margaret L. Schlichting

## Abstract

Adults are capable of either differentiating or integrating similar events in memory based on which representations are optimal for a given situation. Yet how children represent related memories remains unknown. Here, children (7-10 years) and adults formed memories for separate yet overlapping events. We then measured how successfully remembered events were represented and reinstated using functional magnetic resonance imaging (fMRI). We found that children formed differentiated representations in hippocampus—such that related events were stored as less similar to one another compared to unrelated events. Conversely, adults formed integrated representations, wherein related events were stored as more similar, including in medial prefrontal cortex (mPFC). Furthermore, hippocampal differentiation among children and mPFC integration among adults tracked neocortical reinstatement of the specific features associated with the individual events. Together, these findings reveal that the same memory behaviors are supported by different underlying representations across development. Specifically, whereas differentiation underlies memory organization and retrieval in childhood, integration exhibits a protracted developmental trajectory.

## Introduction

In everyday and educational contexts alike, we regularly encounter overlapping information. In the adult brain, such overlap can yield representations for related events that are alternately less similar, i.e., differentiated, (Chanales, Oza, Favila, & Kuhl, 2017; Favila, Chanales, & Kuhl, 2016; Hulbert & Norman, 2015) or more similar, i.e., integrated, (Collin, Milivojevic, & Doeller, 2015; Molitor, Sherrill, Morton, Miller, & Preston, 2021; Schlichting, Mumford, & Preston, 2015) to one another relative to unrelated events. While these discrepant differentiated and integrated traces may both enable later neocortical reinstatement of event features, they may ultimately support different behaviors depending on the task demands: namely memory specificity and flexibility, respectively. Yet, and despite much theoretical interest in this topic (Bauer & Varga, 2017; Keresztes, Ngo, Lindenberger, Werkle-Bergner, & Newcombe, 2018), how children represent related experiences to enable successful memory reinstatement and memory-guided behavior remains unknown. Here, we measure such representations directly to test the hypothesis that children and adults achieve memory success through fundamentally different representational schemes due to ongoing maturation of the hippocampus and medial prefrontal cortex (mPFC) across childhood and adolescence (Bauer, Dugan, Varga, & Riggins, 2019; Ghetti & Bunge, 2012; Keresztes et al., 2018; Østby, Tamnes, Fjell, Westlye, Due-Tønnessen, & Walhovd, 2009; Schlichting, Guarino, Schapiro, Turk-Browne, & Preston, 2017). As children’s abilities to successfully encode and retrieve memories for related events predicts flexible memory behavior (Bauer, & San Souci, 2010) and in turn academic success (Varga, Esposito, & Bauer, 2019), isolating the representations that children leverage during successful memory retrieval is key.

While children can successfully retrieve and make flexible decisions about overlapping experiences (Bauer & San Souci, 2010; Ngo, Newcombe, & Olson, 2018; Rollins & Cloude, 2018; Schlichting et al., 2017), the neural organizational scheme that supports these behaviors has not been identified. For example, children can draw novel connections across memories to make inferences (Schlichting et al., 2017; Shing, Finke, Hoffmann, Pajkert, Heekeren, & Ploner, 2019) and generate new knowledge (Bauer & San Souci, 2010; Bauer, Cronin-Golomb, Porter, Jaganjac & Miller, 2021) at above-chance levels, albeit not as well as adults (Bauer et al., 2021; Schlichting et al., 2017; Shing et al., 2019). However, behavioral success on such tasks should not be taken as evidence of any particular neural organizational scheme, as it can be supported by a variety of different underlying representational codes (Kumaran & McClelland, 2012; Preston & Eichenbaum, 2013; Schlichting & Preston, 2015; Varga, Morton, & Preston, 2024). For instance, inference behavior may be underpinned by either directly accessing integrated neural representations (Molitor et al., 2021; Schlichting et al., 2015; Schlichting & Preston, 2014; Varga & Manns, 2021; Zeithamova, Dominick, & Preston, 2012) or recombining separate memories as needed (Bauer, Varga, King, Nolen & White, 2015; Schlichting, Guarino, Roome, & Preston, 2022; Wilson & Bauer, 2021).

Moreover, integration in the adult brain arises during learning, even in the absence of any requirement to make inferences (Molitor et al., 2021; Schlichting et al., 2015; Schlichting & Preston, 2014; Varga & Manns, 2021; Zeithamova et al., 2012).

Such evidence indicates that neural coding schemes are a means through which adults sort overlapping information, further highlighting the distinction between underlying neural representation and overt behavior. Current developmental theories stemming from behavioral work alternately suggest that children rely more on either representation of commonalities (Keresztes et al., 2018) or storage of distinct memory traces (Bauer & Varga, 2017; Schlichting et al., 2022) to represent related events. The coexistence of such perspectives underscores the challenges associated with drawing conclusions about representation from behavior alone, as well as the need to measure underlying neural representations directly. Here we address this challenge, testing whether integrated or differentiated neural organization supports overlapping memory in children.

We hypothesize that hippocampus and mPFC may be the source of developmental differences in neural organization of highly related events. Hippocampus has been shown to store both integrated and differentiated traces in adults, flexibly aligning representations based on task demands or goals (Chanales et al., 2017; Collin et al., 2015; Favila et al., 2016; Hulbert & Norman, 2015; Molitor et al., 2021; Schlichting et al., 2015). Prior adult data further implicates mPFC in integration of related events (Preston & Eichenbaum, 2013; Schlichting & Preston, 2015; Varga et al., 2024; Zeithamova et al., 2012), showing that mPFC represents connections across learning experiences regardless of the surface features of the task (Morton, Schlichting, & Preston, 2020; Schlichting et al., 2015). Consistent with the notion that memory development may entail a shift from greater reliance on hippocampus in childhood to mPFC in adulthood (Brod, Lindenberger & Shing, 2017), children have been shown to engage hippocampus while retrieving related experiences (Sastre, Wendelken, Lee, Bunge & Ghetti, 2016). Specifically, children show greater hippocampal activation when they successfully recall the details of specific yet overlapping events, relative to when they are unsuccessful (Sastre et al., 2016). The developing hippocampus may thus show a bias toward disambiguating—or differentiating—related events earlier in life; a neural organizational scheme that may be especially important in childhood as a means of minimizing interference between highly similar events (Darby & Sloutsky, 2015).

However, these past findings cannot tell us how related experiences are organized in hippocampus and mPFC, nor whether there are developmental differences in the neural organization that supports successful retrieval. Asking this question requires going beyond measuring whether a region is more or less active at different ages, but rather requires quantifying the similarity of representational patterns for related events (Callaghan et al., 2021; Fandakova, Leckey, Driver, Bunge & Ghetti, 2019; Kazemi, Coughlin, DeMaster & Ghetti, 2022; Qin, Cho, Chen, Rosenberg-Lee, Geary & Menon, 2014), which has yet to be done in a developmental memory investigation in either hippocampus or mPFC. Given that PFC (Fandakova, Selmeczy, Leckey, Grimm, Wendelken, Bunge & Ghetti, 2017; Gogtay et al., 2004; Ofen, Kao, Sokol-Hessner, Kim, Whitfield-Gabrieli & Gabrieli, 2007; Østby et al., 2009) and its connectivity to hippocampus (Calabro, Murty, Jalbrzikowski, Tervo-Clemmens, & Luna, 2020; Simmonds, Hallquist, Asato & Luna, 2014) continues to mature through the third decade of life (Murty, Calabro, & Luna, 2016; for review), we anticipated that integration mechanisms supported by this region would be late to emerge. Specifically, we predicted that adults, but not children, would show memory integration in mPFC. We further predicted that children would instead form differentiated representations for related events in hippocampus, due to earlier maturity of this region relative to mPFC (Brod et al., 2016; Murty et al., 2016), and extant work showing that children engage hippocampus during successful overlapping memory retrieval (Sastre et al., 2016).

Consistent with these predictions, one recent study suggests that younger participants are less likely to form integrated memories relative to adults (Schlichting et al., 2022). That study examined how participants reactivated a previous memory as they encoded a new, overlapping event. While adults reactivated related memories during encoding, in younger participants such reactivation was reduced or even absent. This study implies a developmental shift from storing related memories in non-overlapping neural representations to increasingly overlapping—or integrated—representations, which further complements recent data showing that hippocampal differentiation is at least partially operational in childhood (Benear et al., 2022). However, neither of these studies measured neural representational schemes for related events that were newly learned and subsequently successfully remembered, and therefore cannot address how overlapping events are organized in the developing brain to support memory behavior. Here, we measure the organizational schemes in hippocampus and mPFC for newly formed memories in children and adults as they make memory-based decisions, and how those representations relate to successful behavior at different ages.

We also interrogate the relationship between neural organization and neocortical reinstatement of associated memory features. In addition to organizing related events through differentiation or integration, hippocampal representations also bind distributed neocortical traces together, enabling high-fidelity reinstatement of individual event features at retrieval (Marr, 1971). Similarly, the mature mPFC is thought to combine distributed cortical input into situational models of the environment (Chan, Niv & Norman, 2016; Schuck, Cai, Wilson & Niv, 2016; Wikenheiser & Schoenbaum, 2016), sometimes referred to as memory schemas (Varga et al., 2024), that can later be retrieved to support both new learning and flexible decision making (Preston & Eichenbaum, 2013). The way in which both hippocampal and mPFC representations are organized may thus have downstream consequences for neocortical reinstatement, and in turn, memory behavior.

Past work has shown that hippocampus plays a key role in guiding neocortical reinstatement in adults, with hippocampal BOLD activation tracking the degree to which memory elements are reinstated in neocortex (Ritchey, Wing, LaBar & Cabeza, 2013; Trelle et al., 2020). Stronger neocortical reinstatement, in turn, has been associated with successful and faster memory decisions (Kuhl & Chun, 2014; Mack & Preston, 2016). Here, we ask whether memory reinstatement in higher-order perceptual regions, namely ventral temporal cortex (VTC) and parietal cortex (Kuhl & Chun, 2014; Trelle et al., 2020), predicts upcoming mnemonic decision success in children as it does in adults (Mack & Preston, 2016). Furthermore, we interrogate whether this degree of neocortical reinstatement tracks how related events are represented relative to one another in hippocampus and mPFC. We test whether different representational schemes— differentiation and integration–lead to high-fidelity neocortical reinstatement in children and adults, respectively. Linking successful memory reinstatement to distinct neural organizational schemes in children and adults has important implications not just for foundational theories of memory development (Reyna & Brainerd, 2011), but also for educational practices that seek to maximize memory retention and flexibility at different ages (Brod, 2021).

In the present study, children and adults learned to associate different objects with the same person or place. To ask how those memories were represented, we leveraged high-resolution fMRI and representational similarity analysis (RSA; Kriegeskorte, Mur & Bandettini, 2008) to compare neural activation patterns for objects that were related through a common person or place versus an unrelated baseline. We tested whether children form either differentiated or integrated traces in hippocampus and mPFC, adjudicating alternate perspectives stemming from the cognitive developmental literature (Bauer & Varga, 2017; Keresztes et al., 2018). We further tested whether these hippocampal and mPFC organizational schemes were related to neocortical reinstatement by using multi-voxel pattern analysis (MVPA; Norman, Polyn, Detre, & Haxby, 2006) and RSA to measure reactivation of the associated person/place. Our goal was to test whether differentiation in children’s hippocampus tracks reinstatement on a trial-by-trial basis, and whether the same is true in adults for integration in mPFC. To this end, we focused on middle childhood (7-10 years), as children this age can retrieve associative features (Riggins, 2014) and overlapping (Bauer & San Souci, 2010) events. Although such behavioral evidence suggests that the developing brain can support successful memory retrieval, here we ask a more detailed question: Namely, whether even similar memory behaviors in children and adults might be underpinned by either different hippocampal or mPFC memory representations, and/or different capacities for high-fidelity neocortical reinstatement.

## Methods

### Participants

The current report focuses on MRI data collected from 52 individuals recruited from the greater Austin, TX area: 25 adults aged 19-30 years (*M* = 22.32; *SD* = 3.29; 14 self-reported as female and 11 as male) and 27 children aged 7-10 years (*M* = 9.10, *SD* = 1.09; 15 females and 12 males, based on parental report). Eighty volunteers (*Range* = 7.16-30.09 years) participated in the initial behavioral screening session. During that session, participants (or their parents) completed a series of measures to confirm that participants met screening criteria for the MRI study, which assessed: (1) basic participant information (self- or parent-reported), including right handedness, no color blindness, and whether English was a native language, (2) general intelligence in the normal range or higher (self-completed), as indexed by the vocabulary and matrix reasoning subtests of the Wechsler Abbreviated Scale of Intelligence, Second Edition (WASI-II; Wechsler, 2011; inclusion threshold: <2 standard deviations below the mean), and (3) psychiatric symptoms below the clinical range (self- or parental-reported), as indexed by the Symptom Checklist (SCL)-90-Revised (Derogatis, 1977) in adults (inclusion threshold: standardized global severity score <= 62) and the Child Behavior Checklist (CBCL; Achenbach, 1991) in children (inclusion threshold: standardized global severity score <= 63).

Of the 80 volunteers who participated in the initial screening session, 17 were excluded before the MRI scanning session due to voluntary withdrawal owing to discomfort in the mock scanner (*N* = 2 children), scheduling conflicts (*N* = 2 children and 1 adult), neuropsychological assessments in the clinical range (*N* = 2 children and 4 adults), MRI contraindications (*N* = 2 children and 1 adult), left-handedness (*N* = 1 child), or failure to identify the requisite number of familiar stimuli for the learning task (*N* = 2 children). All participants met our IQ inclusion threshold. Thus, 63 of the 80 participants who were screened returned for the MRI session.

Of the 63 participants who participated in the MRI session, 11 children were excluded due to: (1) low behavioral performance (<90% final learning after five runs [*N* = 1] or failing to perform significantly above chance [50%] on the scanned recall task [*N* = 3]), (2) excessive motion during the functional MRI scans (*N* = 1; see MRI image processing below), or (3) incomplete data (*N* = 6). Incomplete data was operationalized as failure to provide all three runs of the scanned recall task or at least two (out of three) runs of the category localizer task. Because a primary aim was to measure representation of overlapping memories—through comparing similarity of the three object cues that shared the same retrieval target, each of which was presented in a different run of the recall task—loss of a single recall run precluded full assessment of the similarity among the related items from a corresponding set. Moreover, to ensure that the trained MVPA classifier could accurately detect when participants were viewing faces and scenes in the localizer task (see below), at least two runs were needed, one of which was used to train the classifier and one of which was used to test the trained algorithm. There were no issues with data loss for the item localizer task, thus precluding any need for exclusion.

Our target sample size in each age group (*N* = 25) was guided by prior, related research in adults (Mack & Preston, 2016), which showed that this number was sufficient to detect reliable item- and category-level reinstatement in hippocampus and neocortex, as well as hippocampal-neocortical correspondences. Our combined child and adult sample size also aligns well with past developmental work employing multivariate representational methods similar to those used here (Callaghan et al., 2021). An approximately equal number of children were recruited of each age within the 7-10 range and careful efforts were made to achieve approximately equal numbers of males and females within each age. The final sample consisted of native English speakers who identified as 2% American Indian or Alaskan (*N* = 1 child), 15% Asian (*N* = 2 children and 6 adults), 4% Black or African American (*N* = 1 child and 1 adult), 73% White (*N* = 21 children and 17 adults), and 6% mixed racial descent (*N* = 2 children and 1 adult). Twenty-one percent of the sample was Hispanic (*N* = 6 children and 5 adults).

All participants and parents/guardians (hereafter referred to as “parents”) provided verbal assent at the start of the screening session as well as written informed consent at the start of the MRI session. Participants received monetary compensation at each session, at a rate of $10/hour for the screening session and $25/hour for the MRI session. There was also an opportunity to obtain up to a $10 bonus during the MRI session, based on the number of runs completed. All procedures were approved by the University of Texas at Austin Institutional Review Board.

### Experimental Approach Overview

The overarching goal of the present research was to test what representational scheme supports overlapping memory retrieval within the developing brain. Therefore, our priority for the experimental approach was to select a task and design that, should children have access to either differentiated or integrated neural codes, we would be able to detect it. To that end, we adapted a well-established developmental memory task (e.g., Fandakova et al., 2019; Sastre et al., 2016), whereby children and adults were presented with novel face-object and scene-object associative pairs and tested for memory of the individual associations. Critically, each face and scene stimulus was paired with three different objects (e.g., face1 - object1, face1 - object2, and face1 - object3), forming sets of overlapping pairs. Thus, although objects were never directly paired together, they were related through a shared face or scene, thereby allowing us to ask how related objects were organized relative to one another (i.e., differentiated vs. integrated).

Because one goal was to measure differentiated and/or integrated organization of related memories in the developing hippocampus, we chose to address this question with a task that has previously demonstrated the role of hippocampus during overlapping memory retrieval in children (Sastre et al., 2016) and that has also been used to quantify hippocampal and mPFC organization in adults (e.g., Molitor et al., 2021; Schlichting et al., 2015). A primary study that guided our task choice (Sastre et al., 2016) used a similar method to ours, in which children and adults learned multiple objects (16 in the prior work) paired with the same scene. The study showed that hippocampal activation tracked successful overlapping memory retrieval in children; however, only in high performers. Thus, while we could have used other tasks to assess our questions about neural organization of overlapping events (e.g., Bauer & San Souci, 2010; Ngo et al., 2018), we chose the current task because there was prior univariate evidence to suggest that it was sensitive hippocampal recruitment, our *a priori* region of interest (ROI).

Importantly, the performance-related finding from Sastre and colleagues (2016) further suggests that hippocampal recruitment—or hippocampal neural organizational representations, in our case—might only be detected in instances when children successfully form and retrieve memories for the overlapping events. This finding thus informed our second design choice which aimed at producing high memory performance in children. Specifically, in contrast to the prior work that used 16 overlapping objects with the same scene (Sastre et al., 2016), here we limited the number of objects to only 3 per each of the 12 overlapping faces/scenes (i.e., 12 overlapping sets; 36 individual pairs). We know that overlapping learning is challenging for children (Schlichting et al., 2017) and becomes increasingly difficult as the number of overlapping items increases (i.e., 4 vs. 2; Bauer & Larkina, 2017). Thus, by reducing the number of objects that shared an overlapping element, we reasoned that both low- and high-performing children should be able to learn and remember the objects relative to one another; either through differentiating them into disambiguated representations or integrating them within the same representation. Indeed, to foreshadow the results, we showed that 37 out of 38 children learned the pairs, suggesting that the neural organizational patterns reported in the present report are likely to reflect representational capacities available to most children at this age.

In addition to building on well-established developmental tasks and implementing overlapping learning set sizes that were intended to facilitate success, it was also necessary to choose a task that can produce both differentiation and integration neural coding schemes, particularly one that has replicated this finding in the literature in the context of small trial counts. Thus, our third reason for choosing the present approach is that past adult work with this task has shown that 12 overlapping items are sufficient to detect integration and differentiation, in hippocampus as well as in mPFC (Molitor et al., 2021; Schlichting et al., 2015). Based on these neural organization findings in adults, together with past evidence for hippocampal involvement during successful overlapping memory retrieval in children (Sastre et al., 2016), we reasoned that this experimental approach was well-suited to detecting the neural organizational scheme(s) that support overlapping learning and memory retrieval in development.

Finally, although past work indicates that adults have access to both differentiated and integrated organizational schemes, given that our current design was geared toward making the learning task developmentally feasible, we did not necessarily expect adults to show differentiation or integration in hippocampus in this particular learning situation. Prior work has shown mPFC integration in adults under multiple learning conditions (Schlichting et al., 2015), whereas hippocampus in adults can show evidence for either integration, differentiation, or both (Molitor et al., 2021), which seems to depend upon the specifics of the learning situation (Schlichting et al., 2015). Because our *a priori* hypothesis was that children would not form integrated representations, particularly within mPFC (Brod et al., 2017; Fandakova et al., 2017), we further designed our learning task such that it would allow us to observe mPFC integration in the mature brain. To this end, here the novel pairs were comprised of familiar stimuli for which participants had prior knowledge. Adults have been shown to leverage existing knowledge to scaffold integration of familiar information (Bein, Reggev, & Maril, 2020; van Kesteren, Rijpkema, Ruiter, Morris, & Fernández, 2014); something they are more likely to do than children (Bauer et al., 2021; Schlichting et al., 2022), particularly within mPFC (Brod et al., 2017). Therefore, by utilizing familiar stimuli, we reasoned that our design would enable us to show that, under the same set of learning conditions, adults integrate while children may not.

### Stimuli

Stimuli consisted of color images of six faces (three females; three males), six scenes, and 36 objects. All stimuli were sized to 400x400 pixels. Face and scene stimuli were images of characters and places, respectively, obtained from child-friendly films or television shows. Objects were common, everyday items, not obtained from films or television shows. Each face and scene was randomly paired with three different objects, for a total of 36 pairs comprising 12 overlapping sets. This structure allowed us to ask how individual objects that were associated with the same face or scene were represented with respect to one another, as well as how they could each give rise to reinstatement of a neural pattern representing a particular face or scene.

Because developmental differences in neural organizational schemes could result from differences in the ability to retrieve the faces or scenes associated with the object cues, we used familiar stimuli curated to each individual participant. More specifically, only stimuli that participants rated as highly familiar and could verbally label were included in their stimulus set. Furthermore, as discussed above, we hypothesized that adults, but not children, would integrate overlapping stimulus pairs. As such, the use of familiar stimuli also served to test whether adults would leverage existing knowledge to scaffold integration of the new pairs (Bein et al., 2020; van Kesteren et al., 2014), something we did not expect children would do (Bauer et al., 2021).

### Procedure

The experiment consisted of two sessions: (1) an initial behavioral screening session when participant interest in and eligibility for the MRI scanning session were assessed and (2) an MRI scanning session when all the main behavioral and imaging data were acquired. The delay between sessions ranged from 1 to 168 days (*Mean* = 34.17; *SD* = 29.69; *Median* = 27.0). Notably, the delay never exceeded six months, thus ensuring that standardized assessments used to determine eligibility remained valid indicators of neuropsychological and cognitive function over the course of one’s participation. Moreover, task instructions were reviewed at the start of the MRI session, removing any memory burden and the need to control delay intervals.

#### Behavioral screening (Session 1)

To gauge interest and comfort with the MRI protocol that would be implemented at Session 2, participants (and their parents, if minors) were guided through the procedures to anticipate during the MRI session, including the imaging and behavioral task protocols. Eligibility for the MRI session was assessed through standardized and experiment-specific tasks. The screening session had four phases: mock scanner practice, standardized assessments, stimulus familiarity rating task, and behavioral practice tasks.

##### Mock scanner practice

Following informed assent and consent regarding the screening procedures, participants were exposed to the mock MRI scanner where they practiced lying supine in the bore for 2 minutes while audio recordings of scanner noises were played aloud.

##### Standardized assessments

Following the mock scanner, participants and/or their parents completed a series of general intelligence and neuropsychological measures to confirm that participants met screening criteria for the MRI (i.e., WASI-II and CBCL/SCL; see above).

##### Stimulus familiarity rating task

To encourage robust recall (and integration, if adults) and to control for the potential influence of familiarity on neural reinstatement, a customized stimulus set was generated for each participant. Participants were shown a maximum of 326 images (67 faces, 99 scenes, and 160 objects), presented randomly one at a time on a computer screen. Participants were instructed to indicate how familiar each image was, by choosing one of three options: (1) not at all, (2) a little bit, or (3) very. For images judged as “very” familiar, participants were then asked to name or describe the image, which allowed us to control for differences in verbally expressible knowledge. Participants made their responses aloud and had an unlimited amount of time to do so. Only images that were judged as highly familiar and correctly labeled were included in the stimulus set. This and all other experimental tasks were designed and administered using custom scripts in MATLAB R2017a (MathWorks, Natick, MA), which enabled automatic termination of the task when the number of images needed for the primary learning task was obtained. Thus, most participants did not provide familiarity ratings for all 326 stimuli.

Although stimulus sets necessarily differed across participants, several steps were taken to match the types of stimulus content included across participants’ stimulus sets. First, only three object images from a semantic category (e.g., foods, athletic equipment, musical instruments, etc.) could be included in the same stimulus set.

Second, and similar to the object inclusion criteria, we ensured that each of the faces and scenes were drawn from distinctive categories. That is, only one face or scene image from a particular film or television show could be included in the same stimulus set. Together, these criteria served to ensure that all stimulus sets spanned a variety contexts and categories and had matching levels of interfering items.

##### Behavioral practice tasks

Finally, participants practiced short versions of the four experimental tasks that would be performed during the MRI session (see below), ensuring that participants could perform the tasks prior to the MRI scanning visit.

Participants repeated each task until a performance criterion of 90% was reached, which was typically achieved on the first round. The practice task stimuli did not overlap with those used in the main MRI experiment.

#### MRI scanning (Session 2)

Participants/parents provided informed consent and were then reminded of the task instructions and repeated the same behavioral practice tasks as those completed at Session 1. Following the practice tasks, participants then completed the following four experimental phases: pre-exposure item localizer (arrow detection task; in the MRI), pair learning (initial learning/retrieval task; outside of the MRI), memory recall (match/mismatch decision task; in the MRI), and category localizer (repeat detection task; in the MRI).

##### Pre-exposure item localizer (arrow detection task)

The fMRI data acquired during the pre-exposure phase was used to estimate patterns of neural activity associated with perception of the face and scene items that would later be retrieved during the scanned memory recall task, thus enabling quantification of the extent to which these item patterns were reinstated. Participants were exposed to the six face and scene images that comprised the to-be-learned object-face and object-scene pairs, presented through a rapid event-related design. Critically, the pre-exposure phase occurred prior to pair learning, thus allowing for estimation of the neural patterns associated with each item, prior to forming an association with the object cues.

On each trial, a face or scene stimulus was presented for 1.5s with a black fixation dot superimposed on the center of the image. After a random delay (250-750 ms) from stimulus onset, the fixation dot changed to a left- or right-facing arrow. The arrow was presented for the remainder of the 1.5s stimulus presentation and for an additional 1s after stimulus offset. Participants were instructed to respond as quickly as possible to the arrow direction by pressing one of two buttons on an MRI-compatible button box. The concurrent arrow task was intended to ensure attention to the stimuli.

Following the offset of the arrow, a black fixation dot was presented alone for 2.0, 3.5, or 5.0s, with each inter-trial interval (ITI) randomly jittered but occurring approximately equally often for an average ITI of 3.5s across the total run. Each face and scene was presented three times in a pseudo-random order, such that at least two intervening images were presented in between repetitions of the same face or scene image. The number and order of left and right arrows was randomized. Participants completed three consecutive runs, each lasting 4.8 minutes, amounting to a total pre-exposure localizer phase of about 15-20 minutes when accounting for brief breaks and check-ins in between each run.

Task performance was defined as the proportion of arrows for which the arrow direction was correctly detected, averaged across runs. Performance was high in both children (*M = 0.98; SD = 0.03*; *Range* = 0.89-1.00) and adults (*M = 0.99; SD = 0.01*; *Range* = 0.97-1.00), indicating that participants of all ages attended to the images during the pre-exposure item localizer scans.

##### Pair learning (initial learning/retrieval task)

Before the scanned memory recall task, participants completed the pair learning task (Figure 1a) outside of the MRI scanner, which was intended to encourage the formation of strong memories, as evidenced by successful retrieval of the target faces and scenes associated with each object on a three-alternative forced-choice memory test. Participants performed several consecutive learning-retrieval blocks to a 90% criterion. On each trial of the learning block, a face-object or scene-object pair was presented for 3.0s followed by a 0.5s fixation cross (Figure 1a). The face and scene images appeared on the left side of the screen while the object images appeared on the right side of the screen. Each of the 36 pairs was shown once within the learning block, in a random order. Participants were instructed to view the pairs and to create a story relating the two images together.

**Figure 1.**
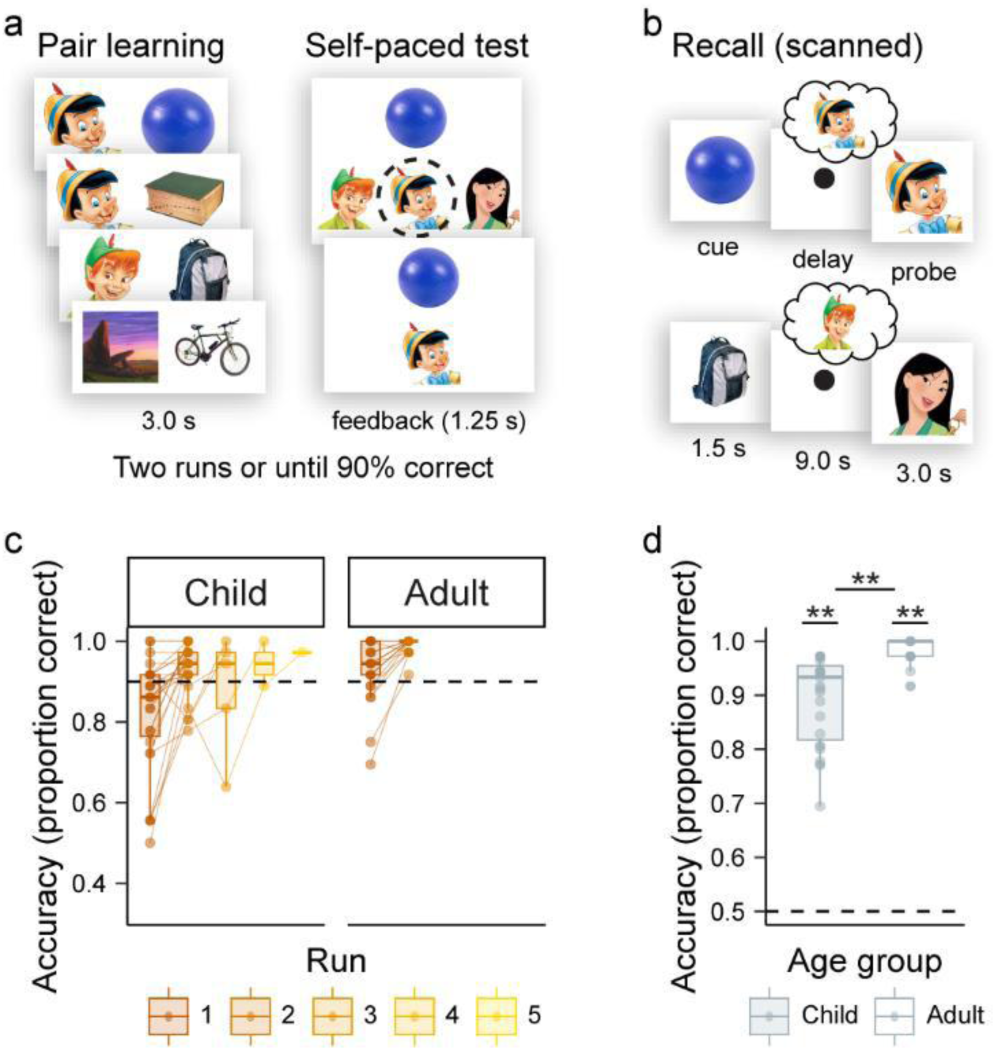
Procedure schematics and corresponding performance on the behavioral tasks. **a** Initial learning/retrieval task. On each run, participants studied the same 36 person-object and place-object pairs and then completed a self-paced three-alternative forced choice memory test with corrective feedback. **b** Scanned recall task. High-resolution fMRI activity was measured while participants viewed previously learned objects and retrieved the target associates (person or place). Associative memory recall was considered successful based on performance on a subsequent mnemonic decision task, in which participants made a button-press response to indicate whether a same-category memory probe (person or place) matched or mismatched the item retrieved. We measured the degree to which neural representations differentiated or integrated overlapping memories during the object cue period, as well as the specificity and degree of ensuing neocortical reinstatement of the associated elements during the delay period. **c** Initial learning/retrieval accuracy. The proportion of correctly selected forced-choice associates was averaged across trials, separately by age group. Dots reflect individual participant means. The lines connecting dots across runs depict within-participant learning trajectories. The number of participants varied by run, depending on how many learning/retrieval repetitions were required to reach the 90% learning criterion (depicted by the dashed black line): Runs 1 and 2: n = 52 (27 children; 25 adults); Run 3: n = 5 children; Run 4: n = 2 children, Run 5: n = 1 child. **d** Scanned recall accuracy. The proportion of correct match/mismatch decisions was averaged across runs and participants, separately for each age group. Dots reflect individual participant means. The dashed black line reflects chance-level performance (0.50). Asterisks above each bar reflect significant above-chance performance at the individual group level while asterisks between bars indicate a significant group difference at a threshold of *p* < .001 (**). For **c** and **d**, box plots depict the median (middle line), 25^th^ and 75^th^ percentiles (the box), and the largest individual participant means no greater than the 5^th^ and 95^th^ percentiles (the whiskers). Individual participant means that extend beyond the whiskers are considered outliers, defined as values that were 1.5 times greater than the interquartile range (IQR).

Following the learning portion, participants were tested on their memory for the 36 pairs. On each trial, an object cue appeared above three face or scene options, one of which was the correct associate (Figure 1a). Participants were instructed to choose the face or scene image that was originally paired with the object by pressing one of three buttons on the computer keyboard (self-paced). Distracter options were pseudo-randomly assigned, with the constraint that foils were always other items from the learning set drawn from the same visual category as the target item. That is, if the target answer was a face, both distracters were also faces. This design criteria served to ensure that participants were able to retrieve the specific association, rather than the gist of the category. Following each choice, regardless of accuracy, a feedback screen appeared for 1.25s, which featured the object together with the correct face/scene. The feedback screen and subsequent test trial were separated by a 0.5s fixation screen.

The pairs were tested in a random order and the correct answers were randomly distributed across the first, second, and third position options. Participants performed at least two runs of learning-retrieval blocks, with different randomized trial orders across each of the runs. If the test accuracy was greater than or equal to 90% on the second run, participants continued on to the scanned memory recall task. Otherwise, participants repeated the learning-retrieval blocks until the 90% criterion was reached (maximum repetitions = 5. Notably, only one child was excluded due to failure to reach the accuracy criterion after five learning-retrieval repetitions.

##### Memory recall (match/mismatch decision task)

The fMRI data acquired during the memory recall task was used to test the primary questions regarding how related memories are organized in different age groups, as well as the degree to which reinstatement of associated memory elements is evidenced in hippocampus and neocortex. On each trial of the recall task, the object cue was presented for 1.5s followed by a 9s delay interval in which only a fixation dot appeared on the screen (Figure 1b). Presenting the objects in isolation enabled us to use this initial cue period of the trial to assess how memories for the related pairs were organized.

During the delay period, participants were instructed to hold the associated face or scene in memory in preparation for an upcoming memory decision. Because the delay period was separated from the object cue and only consisted of a fixation dot, this phase of the trial enabled us to measure reinstatement of the associated face or scene. We implemented a 9-second delay here to ensure that children had sufficient time to reinstate a high-fidelity memory trace. Past developmental electrophysiological work has shown age-related differences in the time course of neural processing during associative memory retrieval (e.g., Bauer, Stevens, Jackson, & San Souci, 2012; Rollins & Riggins, 2017). Moreover, children are less likely to show electrophysiological signatures of memory retrieval in the same recording windows as those observed in adults, but do show similar signatures later in recording windows (Hajcak & Dennis, 2009). As such, here we opted for a relatively long delay, so that even if children were slower to reinstate memories compared with adults, we would still be able to capture it.

To ensure that participants successfully retrieved the associative memories in the MRI scanner, the recall task further included a delayed match to memory decision (Figure 1b). That is, following the delay, a memory probe was presented that was either the correct target associate (i.e., had been paired with the object previously; a match trial) or an incorrect associate (i.e., had been paired with a different object previously; a mismatch trial). The memory probe appeared for 3.0s. Participants were instructed to judge the probe as a match or a mismatch by pressing one of two buttons on the MRI-compatible button box. As was the case during initial learning/retrieval, mismatch probes were always drawn from the same visual category as the target associate, necessitating retrieval of the specific target item rather than gist category information.

Finally, to maximize the efficiency of the slow event-related design and the ensuing neural pattern estimates, null fixation events with durations of 4.5, 6.0, and 7.5s were randomly sampled from a uniform distribution and intermixed between each recall trial.

Notably, during piloting, we observed that children often confused the start and end of each trial. That is, without explicitly prompting for a decision during the probe presentation, children sometimes confused the memory cues (i.e., objects) and probes (i.e., faces/scenes), leading to match/mismatch responses to the object cues rather than the face/scene probes. We took careful steps to eliminate this confusion, as proper identification of the cue periods was necessary to ensure that participants retrieved the target memories. First, the fixation cross that preceded the object cue was presented in green, which was described as a signal that a new trial—and object memory cue—was about to “start”. Second, 1s before the probe presentation elapsed, a red fixation dot was superimposed on the center of the face/scene probe image, signaling that the trial was about to “stop”. Participants were instructed to log a response on the button box when they saw the red dot, if they had not done so already for that memory probe.

Participants completed three runs of the scanned recall task, consisting of 12 trials each. Across the 12 trials within a run, each of the six faces and scenes was a retrieval target once. Moreover, each of the three related objects that shared the same face or scene associate was tested in a different run. Thus, taken together, across the three runs, each of the 36 unique object-face/scene pairs was tested once. Trial order within a run was pseudo-randomized, such that trials from the same stimulus category and condition (e.g., face mismatch trials) never appeared back-to-back. Additionally, target pairs were pseudo-randomly assigned to the match/mismatch condition, with the constraints that (1) within a run, there were three trials of each of the four stimulus category/condition combinations (e.g., face mismatch) and (2) across runs, the same retrieval target (e.g., Pinocchio) was assigned to each match/mismatch condition approximately equally often. Each run lasted 4.10 minutes for a total scanned memory recall phase of about 15 minutes when accounting for brief breaks and check-ins in between runs.

Task performance was defined as the proportion of match/mismatches that were accurately detected across all runs. Above-chance performance was determined for each individual participant using a binomial test of significance (i.e., minimum number = 25/36 trials correct). All participants included in imaging analyses performed significantly above chance on this memory recall task, indicating that participants retrieved the associated memories during fMRI scanning (see Participants above for details on performance-related exclusions). Missing responses during this match/mismatch decision task were rare, yet still occurred (*M*=0.75 trials across participants, *SD*=1.63). To equate the number of trials between analyses that focused on the relationship between neural reinstatement and performance on the match/mismatch decision task, both in terms of (1) accuracy and (2) response time, we opted to exclude missing responses. That is, although missing trials could have been counted as inaccurate responses, because there is no corresponding response time for such trials, including trials without a response would have led to differences in the trial counts between the two analysis types. Thus, for consistency, all reports of match/mismatch accuracy excluded trials for which there was no response (including group means). Importantly, the exclusion of non-response trials did not impact the overall pattern of results nor participant exclusions.

##### Category localizer (Repeat detection task)

At the end of the session, participants performed a blocked, 1-back repeat detection task that included faces, scenes, objects, and scrambled objects. The fMRI data acquired during this task were used to identify functional reinstatement ROIs that were sensitive to face and scene processing (see below; ROI definitions) and to train our MVPA classifier to decode viewing of different stimulus types, thereby enabling estimation of category-level reinstatement during scanned memory recall. Stimuli were drawn from the same set as those used in the stimulus familiarity rating task (see above) but did not overlap with the images selected for the main learning and memory tasks.

Within each run of the repeat detection task, participants viewed a total of 96 images, 24 from each stimulus category. On each trial, a stimulus was presented for 1.5s followed by a 0.5s black fixation dot. Participants were instructed to indicate when a stimulus was identical to (i.e., a repeat of) the immediately preceding image. Stimuli were blocked by category, with six items (including 1 repeat trial) presented per block, lasting a total of 12s. There were four blocks of each stimulus type per run. Stimulus blocks appeared in a pseudo-random order, with the constraint that one of each of the four categories (face, scene, object, scrambled object) occurred before repeating a new set of four category blocks. To ensure that the same stimulus block never appeared consecutively between sets, and to offer participants opportunities to rest periodically within the run, baseline blocks—in which participants viewed a black fixation dot for 12s—were inserted between each set of four category blocks. Participants completed up to three runs of the repeat detection task, with unique stimuli presented in each run.

For some participants, individual localizer runs were not completed due to time constraints. Specifically, 12% of participants (*N* = 6 children) contributed only two of the three possible localizer runs. Each run lasted approximately 4.4 minutes for a total category localizer task of 9-14 minutes, depending on the number of runs completed.

Task performance was defined as the proportion of repeats that were correctly detected, averaged across runs. Performance was high in children (*M = 0.94; SD = 0.07*; *Range* = 0.69-1.00) and adults (*M = 0.98; SD = 0.03*; *Range* = 0.90-1.00), mirroring previous work (Schlichting et al., 2022).

### MR image acquisition

Whole-brain imaging data were acquired on a 3.0T Siemens Skyra system. A high-resolution T1-weighted MPRAGE structural volume (TR = 1.9 s, TE = 2.43 ms, flip angle = 9 degrees, FOV = 256 mm, matrix = 256 x 256, voxel dimensions = 1 mm isotropic) was acquired for co-registration and parcellation. Functional data were acquired using a T2*-weighted multiband accelerated EPI pulse sequence, oriented approximately 20 degrees off the AC-PC axis (TR = 1.5 s, TE = 30 ms, flip angle = 71 degrees, FOV = 220 mm, matrix = 110 x 110, slice thickness = 2 mm isotropic voxels, multiband acceleration factor = 3, GRAPPA factor = 2). At least three fieldmaps were acquired (TR = 647 ms, TE = 5.00/7.46 ms, flip angle = 5 degrees, matrix = 64 x 64, voxel dimensions = 1.7 x 1.7 x 2.0 mm) to correct for magnetic field distortions, which were collected before the first run of each functional MRI task (i.e., pre-exposure item localizer, memory recall, and category localizer) as well as any time the participant exited the scanner for a break in between functional runs within a single functional MRI task.

### MR image preprocessing

Data were preprocessed and analyzed using FSL 5.0.9 (FMRIB’s Software Library, http://www.fmrib.ox.ac.uk/fsl) and Advanced Normalization Tools 2.1.0 (ANTs) (Avants et al., 2011) using a custom preprocessing pipeline (as in Morton, Zippi, Noh, & Preston, 2021). The T1 structural scan for each participant was corrected for bias field using N4BiasFieldCorrection. Freesurfer 6.0.0 was used to automatically segment cortical and subcortical areas based on the processed structural image (Desikan et al., 2006).

Functional scans were corrected for motion through alignment to the center volume using MCFLIRT, with spline interpolation. These realignment parameters were used to compute framewise displacement (FD) and DVARS for each volume. Individual volumes with both an FD and DVARS exceeding a threshold of 0.5 mm were identified. In addition to identifying these high-motion, ‘bad’ volumes, we also marked the volumes immediately before and two volumes immediately after each ‘bad’ volume. If more than 1/3 of the volumes were identified as ‘bad’, the scan was excluded from further analyses. As outlined above (see Participants), one child participant was excluded from analyses because more than 1/3 of the timepoints from a single run of the memory recall task exceeded our FD and DVARS threshold. All other participants contributed all three runs of the pre-exposure and memory recall tasks, and at least two runs of the category localizer task.

Functional scans were unwarped using an adapted version of epi_reg from FSL, which uses boundary-based registration (Greve & Fischl, 2009). This initial boundary-based registration was then followed by ANTs which applied intensity-based registration to refine any misalignments between functional scans and the T1 structural image.

Nonbrain tissue was removed from the unwarped functional scans by projecting a brain mask derived from Freesurfer into each participant’s native functional space. Average brain-extracted unwarped functional scans were registered to a single reference scan collected in the middle of the scanning session (i.e., the second memory recall run) using ANTs. After calculating all transformations, motion correction, unwarping, and registration to the reference functional scan were conducted using B-spline interpolation, in two steps—a nonlinear step followed by a linear step—to minimize interpolation. The bias field for the average image was estimated for each scan using N4 bias correction implemented in ANTs and removed by dividing the timeseries by a single estimated bias field image. Functional time series were high-pass filtered (126s FWHM) and smoothed at 4 mm using FSL’s SUSAN tool. As a result of this registration process, all functional and structural data were coregistered in each participant’s native functional space, defined as the participant’s middle functional run. All within-subject analyses were conducted in this native space, whereas group-level analyses were conducted in template MNI space.

### ROI definitions

Memory reinstatement was examined in two *a priori* anatomical ROIs that have been reliably linked to face and scene reinstatement in past research with adults, including VTC (Gordon, Rissman, Kiani, & Wagner, 2014; Trelle et al., 2020), and parietal cortex (Lee & Kuhl, 2016). Neural organization analyses were conducted in *a priori* anatomical hippocampus, which has been linked to related memory retrieval in past research with children (Sastre et al., 2016) as well as through whole-brain searchlight analyses, which also allowed for test of neural schemes in mPFC. Prior adult findings have revealed different representational schemes in different subregions of mPFC (Schlichting et al., 2015). However, mPFC memory function in children has been interrogated in only a few studies at most (Brod et al., 2017; Calabro et al., 2020; Fandakova et al., 2017). Given the lack of clear theories about mPFC subregion development, we thus chose to use whole-brain grey matter searchlights to interrogate where the largest developmental differences in mPFC organization were apparent.

#### Neocortical ROIs (for item and category reinstatement analyses)

Neocortical anatomical ROIs were bilateral and defined in participants’ native functional space based on the automated Freesurfer parcellation. For the parietal mask, the angular gyrus (AG), supramarginal gyrus (SMG), superior parietal lobule (SPL), and intraparietal sulcus (IPS) were summed (Lee & Kuhl, 2016). For the VTC mask, inferior temporal cortex, fusiform gyrus, and parahippocampal gyrus were summed (Trelle et al., 2020). We then further restricted analysis of neocortical reinstatement to a set of voxels that were most activated during face and scene perception during the separate category localizer task (as in Fandakova et al., 2019), thus ensuring that developmental differences in the spatial extent, spatial location, and/or degree of category-selective cortical activation could not account for any developmental differences observed in reinstatement of item or category neural patterns within these regions (Golarai, Liberman, & Grill-Spector, 2015; Meissner et al., 2019; Peelen et al., 2007; Scherf, Behrmann, Humphreys, & Luna, 2007; 2011).

To define the functional reinstatement ROIs for each participant and neocortical region, for each run of the category localizer, data were modeled using a general linear model (GLM) implemented in FEAT Version 6.00. Stimulus categories (face, scene, object, scrambled object, and fixation) were modeled as five individual regressors. For each category, the four 12.0s-blocks were combined into a single regressor and convolved with the canonical double-gamma hemodynamic response function (HRF). For all task regressors, temporal derivatives were included and temporal filtering was applied. Motion parameters calculated through MCFLIRT and their temporal derivatives were added as additional confound regressors. We separately estimated the contrasts of face above implicit baseline and scene above implicit baseline, which yielded two whole-brain statistical images quantifying voxel-wise activation (one for faces and one for scenes), separately for each run and participant.

The resulting statistical images were averaged across localizer runs for each participant using fixed effects (higher level analysis in FEAT). This yielded two final z-stat images per participant, with the statistical images reflecting how much each voxel was activated by face and scene processing, across all runs. Because functional data were already co-registered across runs, no additional registration step was necessary. Moreover, as the functional reinstatement ROIs were used only for analyses conducted in each participant’s native functional space, no averaging across participants or spatial normalization to a group template was necessary.

The resulting face and scene whole-brain maps were then each masked with the VTC and parietal cortex anatomical ROIs separately. Within each anatomically-masked statistical image, the face and scene ROIs were defined as the 250 voxels showing the greatest amount of activation (i.e., the highest voxel-wise z-stats). We combined the most active face and scene voxels within VTC and parietal cortex separately, yielding two functional reinstatement ROIs for each participant, each containing approximately 500 voxels. Based on prior work showing that face-(Cohen et al., 2019; Golarai et al., 2015) and scene-(Fandakova et al., 2019; Golarai, Liberman, Yoon, Grill-Spector, 2010) representations are highly similar in children and adults when distributed activation patterns are examined, we did not assume spatial contiguity between voxels. Occasionally, the same voxel was identified as among both the most active face and scene voxels, resulting in fewer than 500 voxels (VTC *Range*: Adult = 344-448 and Child = 320-457; Parietal *Range*: Adult = 304-431 and Child = 288-410).

These more specialized functional VTC and parietal masks that contained only the voxels that were most sensitive to face and scene processing for each participant were used to examine both category- and item-level reinstatement, averaging reinstatement evidence across the entire ROI. We also examined item reinstatement via searchlight analyses conducted across the whole-brain grey matter masks (see below), small-volume corrected within each functional reinstatement ROI as well as within anatomical hippocampus (see below; Hippocampal ROIs), revealing item reinstatement in more localized voxels within these larger ROI masks.

#### Hippocampal ROIs (for both item reinstatement and memory organization analyses)

The hippocampal mask was delineated manually on the 1mm MNI152 template based on a cytoarchitectonic atlas (Öngür et al., 2003). The bilateral hippocampal anatomical ROI was reverse normalized into each participant’s native functional space using ANTs. In addition to testing our *a priori* hypotheses regarding neural organization in children, the anatomical hippocampal ROI was also used in whole-brain searchlight analyses interrogating both memory organization and item-level reinstatement, small-volume corrected in order to identify localized voxels that exhibited significant differentiation/integration, or significant item-level reinstatement.

#### Whole-brain grey matter ROIs (for both item reinstatement and memory organization searchlight analyses)

Whole-brain grey matter ROIs were defined in participants’ native functional space based on the automated Freesurfer parcellation. While searchlight analyses testing for significant item reinstatement and memory organization were conducted within each participant’s individual grey matter mask, for group-level statistical testing, group masks were generated by adding individual participant masks together, normalized to MNI group space using nonlinear SyN transformations in ANTs.

### Deriving event-specific activation patterns (inputs to MVPA and RSA analyses)

Three parallel sets of models were performed to derive event-specific patterns as inputs to our neural reinstatement and memory organization analyses: one model focused on the pre-exposure item localizer task (i.e., item perception) and two focused on the memory recall task (one modeling patterns of activation for each object cue; the other modeling the delay period). For all models, separate event-specific univariate GLMs were conducted for each run, using the least squares single (LSS) method (Mumford, Turner, Ashby, & Poldrack, 2012; Xue, Dong, Chen, Mumford & Poldrack, 2010). Within each run, the individual events were modeled at the trial level across the whole-brain grey matter mask. For the memory recall task, in which a single trial comprised two separate phases of interest (i.e., the object cue and the delay period), separate models were conducted for each. For all models, confound regressors were included to account for motion: six head motion parameters, their temporal derivatives, framewise displacement, and DVARS (Power, Barnes, Snyder, Schlaggar & Petersen, 2012). We also accounted for high-motion ‘bad’ volumes identified during pre-processing, by including additional motion regressors for time points during which head motion exceeded both the FD and DVARS thresholds. All models were conducted on the spatially pre-processed fMRI data. Temporal filtering was further applied to each event during modeling.

#### Modeling item perception (for item reinstatement analyses)

We wanted to quantify the degree of item-specific reinstatement during the delay period of the memory recall task. For this purpose, we estimated neural patterns associated with the perception of each of the six faces and scenes from the pre-exposure item localizer phase. Separate LSS GLMs were conducted for each of the three scanned pre-exposure runs, with each of the six faces and scenes modeled as a single event. That is, each of the three 1.5s stimulus presentations of an individual face/scene stimulus from a run were collapsed into a single regressor and convolved with the canonical double gamma HRF. This approach resulted in one voxelwise beta image for each of the six face and scene images for each run and participant, which were used as inputs to the RSA assessing reinstatement of these item-specific perceptual patterns during recall.

#### Modeling the object cue period (for memory organization analyses)

We modeled the object cue period to interrogate the degree of similarity among patterns of fMRI activation evoked by related versus unrelated objects. To this end, within the memory recall task, we extracted neural patterns for each of the individual object cue presentations, each modeled as a single 1.5s regressor and convolved with the canonical double gamma HRF. Nuisance regressors of no interest that accounted for later trial variance were also modeled, including the delay period (by category: face and scene targets) and probe (category x probe type: face match, face mismatch, scene match, scene mismatch). The resulting beta images consisted of voxelwise parameter estimates for each of the 36 object cue activation patterns (12 per run) for each participant. These voxelwise estimates were used as inputs to the memory organization RSA assessing the similarity of related object cues relative to unrelated object cues.

### Modeling the delay period (for item and category reinstatement analyses)

Finally, we modeled the delay period to quantify the degree of neocortical item- and category-level reinstatement, conducting two separate models that either (1) collapsed activation across the full 9s-delay period or (2) split activation estimates into the first and second halves (4.5s) of the delay period. We extracted neural patterns for the delay period on each recall trial; each modeled as either a single 9s regressor or as two 4.5s regressors and convolved with the canonical double gamma HRF. Additional regressors that accounted for the preceding trial-wise object cue variance were also modeled, as were regressors that accounted for variance during the probe period (category x probe type; as in the object cue model above). The approach resulted in one voxelwise beta image for each of the delay period patterns (36 for the 9s model; 72 for the split-half model) for each participant, which were used as inputs to the MVPA and RSA approaches interrogating category- and item-level reinstatement, respectively.

Notably, in the object cue model above, the events of no-interest (i.e., the delay period) were modeled at the category level (i.e., face and scene targets); whereas, in the delay model here, the events of no-interest (i.e., the object cues) were modeled at the trial level. Our rationale for this difference is that, in the object cue model, the phase of no-interest (i.e., the delay) occurred after the phase of interest (i.e., the object cue), and thus could not be expected to influence the trial-wise estimates of the object cues. Conversely, in the delay model here, the phase of no-interest (i.e., the object cue) occurred before the phase of interest (i.e., delay period), and could therefore carry over into trial-level delay period activation estimates. We thus modeled object cue variance at the trial level to mitigate the influence of any carry-over effects.

### Category-level reinstatement analyses (using MVPA)

The goal of the MVPA classification analysis was to quantify the degree to which category-level information (face vs. scene evidence) was reinstated during the delay period of the memory recall task. Pattern classification analyses were conducted on detrended and z-scored category localizer and recall data at the within-participant level, in each participant’s native functional space separately within each participant’s functionally defined VTC and parietal ROIs (see above) using PyMVPA (Hanke et al., 2009) and custom Python scripts.

#### Training the pattern classifier during category perception

We first ensured that the MVPA classifier could accurately detect when participants were viewing faces and scenes. To this end, a sparse multinomial logistic regression (SMLR) classifier (lambda = 0.1, the default) was trained to discriminate neural activation patterns associated with viewing faces and scenes in the category localizer task. The classification training was conducted on the preprocessed, spatially smoothed functional timeseries data. Each of the individual measurement volumes corresponding to a face or scene block (eight volumes per block) was labeled according to the visually presented category. The fMRI data was shifted by 4.5s to account for the hemodynamic lag and these volume labels were used as inputs to the classifier algorithm for training.

Classification performance was evaluated with a leave-one-run-out cross-validation approach, such that unlabeled fMRI patterns from an individual run (i.e., one “fold” which was “left out” of the classifier training) were labeled as a face or scene, according to the trained classifier from the other runs for that participant. That is, for each unlabeled activation pattern corresponding to an individual volume, the classifier labeled it as either a face or scene—a prediction that was based on the correspondence between the unknown activation pattern and the classifier’s trained algorithm from the other runs. This validation approach was repeated for each run separately, resulting in two or three cross-validation folds, depending on the total number of runs available.

Classifier performance was assessed by averaging the accuracy across the predictions for all cross-validation folds, yielding a single cross-validation accuracy score for each participant and ROI which was compared to chance (50% for two categories).

#### Applying the trained classifier to the delay period recall data

After validating that the classifier could accurately decode patterns of face and scene activation, we then applied the classifier to the memory recall task to quantify the degree to which face and scene patterns were reinstated in memory during the delay period. The same classifier as above was used here, except rather than the leave-one-run-out training approach, here the classifier was trained on all runs of the localizer task for each ROI.

To predict the degree of face and scene evidence during the delay period of each scanned recall trial, the trained classifier was applied to the trial-level delay patterns modeled through the GLM approach (see above). Specifically, for each 9s or 4.5s retrieval pattern (corresponding to the full or split-half delay model, respectively), the pattern-classification algorithm estimated the amount of face and scene activation evidenced out of a probability of 1, yielding two continuous probabilities ranging from 0 (no evidence) to 1 (perfect evidence) for each trial. Chance-level evidence for a target category corresponded to a raw value of 0.5, reflecting equivalent evidence for both categories. To correct for non-normality, the classifier output was transformed to logits (log odds = log[x/(1-x)]) (as in Richter, Chanales, & Kuhl, 2016; Trelle et al., 2020). A logit value of 0 corresponded to a raw, chance-level reinstatement value of 0.5. Larger positive logit values indicated greater evidence for the target category; whereas, negative logit values indicated greater evidence for the nontarget category.

#### Analysis of group-level and trial-level category reinstatement

Reinstatement of the target category was evaluated within and between groups by comparing across-trial mean target reinstatement probability values against chance levels (i.e., 0; one-sample t-tests; one-tailed) and between groups (independent sample t-test; two-tailed). Importantly, we were interested in what *successful* category-level reinstatement looks like and whether such reinstatement differs between age groups.

As such, here analyses were restricted to trials in which performance was successful at both unscanned final learning and scanned memory recall (see below; Performance-related trial restrictions). In addition to group-level analyses, trial-wise estimates for the target category (i.e., face evidence when the retrieval target was a face) were also entered into within-participant mixed effects and drift diffusion models, enabling test of the association between reinstatement and decision accuracy and speed, respectively (see below; Trial-level mixed effects models and Trial-level drift diffusion models, respectively).

### Item-level reinstatement analyses (using RSA)

To characterize the degree to which item-specific information was reinstated during the delay period of the memory recall task, RSA (Kriegeskorte et al., 2008) was used to compute how much more similar delay period retrieval patterns were to perception of the same item (*same-item*) versus to other items from the same category (*different-item*). The RSA was implemented in PyMVPA (Hanke et al., 2009) within functionally defined VTC and parietal cortex (see above) in each participant’s native functional space, as well as across the whole-brain grey matter mask, small-volume corrected within both anatomical hippocampus and functionally defined VTC and parietal cortex (see below; Whole-brain item reinstatement with searchlights). Within each functional reinstatement ROI or searchlight sphere, we computed the similarity between the face and scene perception patterns derived from the pre-exposure item localizer and the retrieval patterns derived from the delay period of the memory recall task, via pairwise Pearson’s correlation coefficients. Differences in pattern similarity between same-item similarity values (e.g., *r* of delayface1 & perceptionface1) and different-item similarity values from the same category (e.g., *r* of delayface1 & perceptionface2) were computed by transforming Pearson’s correlation values to Fisher’s z scores and averaging the similarity values corresponding to the same- and different-item conditions, using custom MATLAB and Python scripts. Because similarity values were always computed between pre-exposure and recall neural patterns, which necessarily occurred across runs, temporal autocorrelation was not a concern (Mumford et al., 2012).

#### Analysis of group-level and trial-level item reinstatement

A mean item reinstatement index was derived for each participant and each functional reinstatement ROI by subtracting the different-item similarity values (always restricted to the same category) from the same-item similarity values, restricted to correct scanned recall and final pair learning trials (see below; Performance-related trial restrictions). These participant-level estimates were averaged across participants within each age group and compared to chance levels (i.e., 0; one-sample t-tests; one-tailed), yielding an estimate of whether item reinstatement was reliably evidenced on successful trials in each neocortical functional ROI. To examine age-related differences, the group-level item reinstatement indices were compared between children and adults (independent samples t-test; two-tailed), separately for each VTC and parietal functional ROI.

In addition to group-level pattern similarity estimates which collapsed across all correct trials and across participants within a group, trial-level item reinstatement estimates were computed at the within-person level, in each neocortical functional ROI separately. Here, we examined whether trial-by-trial variability in delay period item reinstatement corresponded with (1) neural organization of the preceding object cue (see below; Memory organization analyses), and (2) match/mismatch behavior during the subsequent memory probe (see below; Trial-level mixed effects models and drift diffusion models; not restricted to correct trials). For each recall trial, a raw “same-item” reinstatement measure was computed, which compared the similarity between that delay pattern and the same-item perceptual patterns from the pre-exposure phase.

Notably, unlike the group-level analyses in which we tested whether “same-item” reinstatement was reliably observed compared to a “different-item” baseline, here we focus on raw “same-item” neural estimates rather than baseline-subtracted estimates. The latter would have necessitated the use of a different baseline for each individual recall trial. Because varying baselines could differentially influence—and overwhelm— pattern similarity estimates for a singular same-item trial estimate, for all trial-level analyses and results (including those on memory organization; see below) we report raw values without reference to a baseline.

#### Whole-brain item reinstatement with searchlights

A searchlight extension of the main item reinstatement analysis in functionally-defined VTC and parietal cortex was conducted to identify localized voxels that exhibited significant item-level reinstatement, at the whole-brain level, which was then small-volume corrected within the functional ROIs as well as within hippocampus. Here, the same general RSA analysis was repeated, wherein similarity matrices were generated by calculating the pairwise Pearson’s correlation values between the delay patterns and pre-exposure patterns of viewing that same item versus different items, restricted to correct trials and transformed to Fisher’s *z*. Yet rather than calculating one single similarity index for an entire ROI, here the similarity contrasts were computed at the level of individual searchlight spheres (radius = 3 voxels). A random permutation approach was then used to evaluate the statistical significance of the observed index within each searchlight sphere, whereby the actual observed contrast—same minus different item—was compared to a null distribution of indices calculated from randomly permuting the values that contributed to the observed contrast (see below; Statistical analysis of searchlight evidence). To parallel the primary analyses that tested if children and adults show reliable item reinstatement at the group level in VTC or parietal cortex, we tested for significant item reinstatement regions within the child and adult groups separately (see below; Statistical analysis of searchlight evidence).

### Memory organization analyses (using RSA)

To characterize how object cues that shared an overlapping face/scene associate were organized relative to one another, an RSA approach was again implemented in PyMVPA (Hanke et al., 2009). The RSA was conducted in each participant’s native functional space, both within anatomically defined bilateral hippocampus as well as across the whole brain, which further allowed us to interrogate mPFC (see below; Whole-brain memory organization with searchlights). Specifically, within the hippocampal ROI and each whole-brain searchlight sphere, we computed the similarity between the different object neural retrieval patterns derived from the object cue period of the memory recall task via pairwise Pearson’s correlation coefficients.

Differences in pattern similarity between object cues that shared the same retrieval target (i.e., related objects) and objects that shared different retrieval targets from the same category (i.e., unrelated objects) were computed, by transforming Pearson’s correlation values to Fisher’s z and averaging the similarity values corresponding to each relatedness condition via custom MATLAB and python scripts. Only across-run similarity values were included to mitigate the influence of temporal autocorrelation on pattern similarity estimates (Mumford et al., 2012).

#### Analysis of group-level and trial-level memory organization effects

The participant-level related object and unrelated object mean pattern similarity estimates were entered into group analyses, by averaging across participants within an age group. To assess the effect of age group and condition (related vs. unrelated) on neural pattern similarity within our *a priori* hippocampal ROI, a 2 (child, adult) x 2 (related, unrelated) repeated measures analysis of variance (ANOVA) was conducted. Significant interactions and main effects were interrogated through separate repeated measures ANOVAs and/or paired sample t-tests. Chance-level evidence for neural organization corresponded to a nonsignificant condition effect, reflecting that the similarity of neural activation patterns did not differ for related versus unrelated objects. Evidence for neural integration, on the other hand, was indexed by higher pattern similarity for related objects versus unrelated objects. Evidence for neural differentiation was indexed by the opposite pattern, such that neural patterns for related objects was lower than that of unrelated objects. The approach thus enabled us to test whether neural integration and/or differentiation was reliably observed within an age group, and whether these schemes differed by age group.

In addition to group-level pattern similarity analyses which collapsed across all correct trials and across participants in each age group, trial-level neural organization similarity estimates were also computed for each individual participant. As above, for each object cue, the pattern similarity between that object and the two other objects that shared the same face/scene retrieval target was computed and averaged. This trial-level raw “related” neural estimate was then entered into mixed effects models interrogating the association between neural memory organization and ensuing neocortical item- and category-level reinstatement (see below; Trial-level mixed effects models). As for trial-level item reinstatement analyses (see above), here we focused on raw “related” neural estimates rather than on baseline-subtracted neural estimates.

#### Whole-brain memory organization with searchlights

A searchlight extension of the neural memory organization analysis conducted in *a priori* anatomical hippocampus was conducted to identify localized voxels that exhibited significant differentiation or integration of related object cues, both at the whole-brain level— enabling assessment of developmental differences across neocortex, including within mPFC—and within hippocampus. Specifically, we looked for (1) differentiation and (2) integration of related object cues, operationalized as lower or higher pattern similarity for related object cues relative to unrelated object cues (from the same visual category), respectively. The same general RSA analysis as above was repeated here, wherein similarity matrices were generated by calculating the pairwise Pearson’s correlation values between the related versus unrelated object cue patterns, restricted to correct trials and transformed to Fisher’s *z*; calculated within each searchlight sphere (radius = 3 voxels). A random permutation approach was used to evaluate the statistical significance of observed neural organizational effects (see below; Statistical analysis of searchlight evidence).

To evaluate the presence of age-invariant neural organizational effects, we tested for significant neural organization regions collapsed across children and adults. In the integration searchlight, the observed and randomly permuted indices were calculated by subtracting the unrelated similarity values from the related similarity values—in essence, searching for regions that showed related > unrelated within each searchlight sphere. In the differentiation searchlight, we performed the inverse contrast, such that the observed and randomly permuted indices were calculated by subtracting the related similarity values from the unrelated similarity values—thereby reflecting regions that showed unrelated > related within each searchlight sphere.

To test for developmental differences in neural organization, we tested for regions that showed greater neural integration or neural differentiation via two contrasts: (1) adults > children or (2) children > adults. Importantly, an interaction of integration or differentiation with age group is ambiguous with respect to whether significant integration and/or differentiation exists within each age group individually. For example, if some voxels show adults > children for the integration contrast (related > unrelated), this age effect could be attributed to adults exhibiting reliable integration (related > unrelated) while children show either no significant organization (related = unrelated) or differentiation (unrelated > related). Likewise, this effect could also result from adults showing more evidence of integration relative to children, despite no significant evidence of integration (related > unrelated) in the adult group overall. As such, for regions showing a significant effect of age on neural organizational coding scheme, we then separately tested for integration or differentiation within each age group. We report age interactions only when an effect was reliable within at least one group (i.e., enhanced integration was driven by significant integration in one group and/or by significant differentiation in the other group).

### Statistical methods and significance thresholding

#### Performance-related trial restrictions

Where possible, for all statistical analyses that evaluated neural measures of item or category reinstatement or memory organization (i.e., mean-level group analyses, searchlights, and mixed effects models that focused on the relationship between memory organization and reinstatement), we limited analyses to “correct” trials to reduce the possibility that age-related differences in overall performance contaminate our neural results. Specifically, for the aforementioned analyses, we measured both overlapping neural memory organization and item/category reinstatement considering only trials for which the participant both (1) learned the pair initially, which we defined as having remembered it correctly on the final retrieval test from the pair learning phase and (2) correctly responded during the match/mismatch decision of the scanned recall task. The logic is that if participants did not learn the pair initially, or did not correctly identify the probe as a match or mismatch, there is a higher likelihood they were reinstating the wrong memory (or nothing at all).

The performance-related restrictions were applied to all statistical analyses with the exception of two in which we used participant-level category and item reinstatement to predict decision accuracy and speed on the match/mismatch task. In the case of decision accuracy, because we wanted to predict accuracy on the match/mismatch task at the trial level, we needed to include both correct and incorrect trials as outcomes to run our logistic regression. Thus, here, the mixed effects analysis was limited according to neither performance on the final learning task, nor the actual scanned recall task (see below; Trial-level mixed effects models). In the case of response speed, there were no trial restrictions based on behavioral accuracy, as the drift diffusion model (DDM; Ratcliff, 1978) approach requires inclusion of both correct and incorrect trials (see below; Trial-level drift diffusion models).

#### Mean-level group analyses

Group analyses comparing differences between age groups and/or reliable effects within age groups were conducted via t-tests and ANOVAs in SPSS (Version 27), with Bonferroni-adjusted p-values reported to control for multiple comparisons. In cases where the assumption of sphericity was violated in ANOVAs, the degrees of freedom were adjusted using Greenhouse-Geisser corrections. Likewise, for independent samples t-tests, when the Levene’s Test indicated unequal variances between age groups, the degrees of freedom were adjusted accordingly.

Notably, for category and item reinstatement (but not other effects), we evaluated the statistical significance within each group via a one-tailed t-test. The rationale for using one-tailed t-tests here and not for other analyses (e.g., memory organization) is that only for the reinstatement analysis was our prediction strongly directional. That is, we were only interested in effects wherein reinstatement was significantly greater than (not merely different from) zero, and therefore 1-tailed tests are appropriate. By contrast, for memory organization analyses both directions of difference are interpretable (i.e., related < unrelated [less than 0] is taken as evidence of differentiation, whereas related > unrelated [greater than 0] implies integration).

Therefore, throughout the paper all other statistical tests are two-tailed unless otherwise noted.

#### Trial-level mixed effects models

Trial-wise analyses were conducted using mixed effects models using the lme4 package (Bates, Machler, Bolker, & Walker, 2015) in RStudio (2020; version 1.2.5042). Two primary sets of models were performed: (1) one that tested the relationship between item/category reinstatement and match/mismatch behavior and (2) another that tested the relationship between memory organization (integration/differentiation) and item/category reinstatement. For all models, we assessed model fits, first comparing a base model (for which age group was not included as a predictor) to a model that included a main effect of age. We then further compared that best-fit model (Base vs. Main Effect) to a model that included an interaction with age group. Results from the best-fitting model are reported in all cases, with the exception of interaction models, for which we sometimes report interaction models when there is (1) a marginally significant improvement in model fit when including an interaction term, suggesting modest evidence of a conditional effect or (2) a significant interaction term in the model itself, suggesting that this is a reliable effect. For all models, we modeled the participant-specific association as a random slope.

Individual model predictors were scaled and centered across all trials within a participant prior to analyses, thus standardizing the distribution between subjects. For Base and Main Effect models, statistical significance of individual predictors was assessed using a Wald chi-square test (Fox & Weisberg, 2011); whereas, for interaction models, separate slopes were assessed for each group (i.e., simple_slopes function from reghelper R package).

For the model testing whether trial-wise variability in item and category reinstatement explained behavioral performance on the match/mismatch memory task, we conducted a single logistic mixed effect model (glmer function) using a binomial linking function (family = binomial) that included both item reinstatement (same-item similarity) and category reinstatement (logit transformed classifier evidence for target category) indices in the VTC and parietal cortex functional reinstatement ROIs during the delay interval. More specifically, we included all four reinstatement measures—VTC item, VTC category, parietal item, parietal category—in a single model, enabling assessment of which reinstatement predictors (if any) explained unique variance in whether participants answered the subsequent match/mismatch probe correctly. In addition to these central predictors, we also included additional regressors that captured variability in (1) univariate hippocampal activation during the cue period as well as (2) univariate VTC and parietal activation during the delay period, thus mitigating the likelihood that any observed effects of target memory reinstatement on performance are instead due to global changes in BOLD activation. Moreover, as noted above, this analysis did not control for behavioral performance, as this was the dependent measure of interest.

For the models that tested whether hippocampal or mPFC object cue memory organization was associated with neocortical reinstatement, we applied the same performance-related restrictions applied elsewhere (i.e., we restricted to correct final learning and scanned recall trials). Here, we employed linear mixed effects models (lmer function) to test the association between object cue similarity—defined as raw “related” estimates (see above)—separately for anatomical hippocampus and mPFC (searchlight cluster; see Results) and ensuing neocortical reinstatement during the delay period. For the reinstatement dependent measures, we focused on only those ROIs that, in both age groups and across the full delay, either showed (1) reliable reinstatement or for which (2) trial-wise variability in reinstatement was linked to behavior at significant or trend levels. This resulted in three neocortical dependent measures: (1) VTC category, (2) parietal category, and (3) parietal item (see Results).

There was thus a total of six cue organization/reinstatement models, three that looked at the relationship between anatomical hippocampal cue organization with each of the three reinstatement variables and three that looked at the same reinstatement variables, but with mPFC cue organization as the predictor.

In addition to our *a priori* hippocampal and mPFC cue organization regions of interest, we conducted additional follow-up analyses in which we ran the same models for other searchlight regions that showed significant integration in either children or adults. The approach allowed us to test whether the effect of integrated neural organization on neocortical reinstatement were selective to our *a priori* ROIs, or rather observed elsewhere in other regions that showed evidence of this neural coding scheme (either in adults or children). To mitigate the issue of multiple comparisons, here we chose to restrict these follow-up comparisons to only those neocortical dependent variables that showed a significant association with our *a priori* hippocampal and mPFC memory organization variables.

#### Trial-level drift diffusion models

Following previous work (Mack & Preston, 2016), we used DDM (Ratcliff, 1978) to assess whether reinstatement during the delay period predicted, for each individual trial, response speed on the subsequent match decision. In particular, we evaluated the association between drift rate *v*—defined as the rate at which evidence for a memory-based decision accumulates—and response time distributions on the mnemonic decision task. There were no trial restrictions based on behavior applied for the purposes of this analysis, as the DDM approach requires inclusion of both correct and incorrect trials. However, we did restrict our analysis to only *match* trials to equate the difficulty of the decision/matching process, which would be expected to vary across mismatch trials and therefore be an additional source of noise.

Guided by the significant or trend-level associations between VTC category and parietal item reinstatement effects on decision accuracy (see Results), here we constrained our analysis to these same neocortical reinstatement measures, focusing on trial-by-trial evidence of category VTC reinstatement (target reinstatement) as well as item-level parietal cortex reinstatement (raw same item estimates). We modeled the linear effect between VTC category reinstatement and parietal item reinstatement on the drift rate parameter. Within this model, we additionally controlled for mean hippocampal BOLD response on each trial, as this was a significant predictor in the mixed effects accuracy models (unlike BOLD VTC and parietal responses; see Results). The models—each containing three predictors (hippocampal BOLD cue response, VTC category reinstatement, parietal cortex item reinstatement)—were run separately for children and adults, using the Hierarchical Drift Diffusion Model toolbox (HDDM version 0.5.3; Wiecki, Sofer, & Frank, 2013) which implements hierarchical Bayesian parameter estimation using Markov-Chain Montel Carlo (MCMC) sampling methods. Specifically, models were estimated with 5,000 MCMC samples, after discarding the first 1,000 samples for burn-in and thinning of 2, resulting in a distribution of 2,000 regression coefficients.

To evaluate statistical significance among the neural predictors and the drift rate parameter, we computed the proportion of regression coefficients that were more or less than 0 depending on the sign of the coefficient median, constituting a p-value for each neural measure. Only predictors for which the corresponding p-values exceeded the 95^th^ percentile (i.e., p < .05) were considered statistically significant. The models thus resulted in three regression coefficients reflecting the relationship between each of the neural measures (i.e., parietal item reinstatement, VTC category reinstatement, cue-period hippocampal BOLD activity) and drift rate for the match decisions. In situations where reinstatement is high, this should result in a faster accumulation of evidence (or drift rate) and a faster response time. When item reinstatement is low, evidence accumulates gradually and the RT is slower. Therefore, positive coefficients would provide evidence that greater reinstatement is associated with a higher drift rate, or faster evidence accumulation. Notably, all model fits were assessed beforehand, including checks on model convergence using the Gelman-Rubin R-hat statistic (see Mack & Preston, 2016 for details) as well as ensuring that the neural models provided a better fit (i.e., smaller deviance information criterion [DIC] value) than a base DDM model that did not include effects of reinstatement or hippocampal cue activation.

#### Statistical analysis of searchlight evidence

Two searchlight extensions were conducted to identify more localized voxels that exhibited (1) significant item-level reinstatement and (2) significant differentiation and/or integration of related object cues.

All searchlights were run across the whole-brain grey matter masks within each participant’s native functional space. Searchlight maps were then masked and statistically evaluated within each ROI separately. Below we detail the general approach to identifying statistically significant clusters of voxels.

A random permutation approach was used to evaluate the statistical significance of the observed effect within each searchlight sphere, whereby the actual observed contrast—(1) item reinstatement: same minus different item or (2) memory organization: related minus unrelated object cues (for integration; or vice versa for differentiation)— was compared to a null distribution of indices calculated from randomly permuting the values that contributed to the observed contrast. For example, with respect to the item reinstatement searchlight, we randomly shuffled assignment of the same-item and different-item similarity values, resulting in a random item reinstatement index. For each searchlight analysis, the permutation process was conducted 1,000 times within each searchlight sphere, yielding a null distribution of contrast indices. A z-score reflecting how much greater the observed index was compared to the mean of the null distribution was computed, resulting in a whole-brain z-map for each searchlight analysis.

To evaluate whether these voxel-wise z-stats showed statistically reliable neural coding signatures at the group level, we applied a cluster-based random permutation approach. To align individual voxels across participants, we first normalized the participant z-maps to MNI group space using nonlinear SyN transformations in ANTs. Using Randomise in FSL (Winkler, Ridgway, Webster, Smith & Nichols, 2014). We then assessed the mean similarity value across participants, wherein an actual observed value was calculated by averaging the observed z-stat index—reflecting item reinstatement or neural memory organization—across participants for each individual voxel. To evaluate the statistical significance of the observed, group-level index, each across-participant voxelwise z-stat was compared to a null distribution of 5,000 permuted samples via Randomise. The random permutation approach was conducted in one of two ways, depending on the type of group contrast. When evaluating whether a particular group—child, adult, or the full sample (child + adult)—exhibited reliable item reinstatement or neural memory organization, a one-sample t-test was performed. In contrast, when evaluating group differences—children > adults, or vice versa—an unpaired sample t-test was performed.

With respect to the one-sample approach, on each of the 5,000 permutations, the sign of participants’ z-maps was randomly inverted within Randomise. That is, a subset of the participant z-maps were multiplied by -1 and a new, permuted across-participant index was calculated. The 5,000 random inversions were used to compute a voxelwise probability map reflecting the proportion of null distribution sample values greater than or equal to the observed group-level index (i.e., *p* values). Put concretely, a voxel value of p = .01 would indicate that only 50 out of the 5,000 permuted estimates exceeded the actual observed group-level index.

With regard to the two-sample approach, rather than changing the sign of participants’ z-maps, each permuted sample was generated by randomly shuffling assignment of participants to the adult or child group within Randomise. Here, the 5,000 random permutations were used to compute a voxelwise probability map reflecting the proportion of null distribution sample values greater than or equal to the observed group difference, run separately for the adult > child contrast and child > adult contrast. In this case, a voxel value of p = .01 would indicate that only 50 out of the 5,000 permuted estimates exceeded the actual observed group difference.

Finally, following computation of the voxel-wise statistical maps via Randomise, we then sought to identify significant clusters of spatially contiguous voxels (Woo, Krishnan & Wager, 2014), thereby controlling for the family-wise error rate across voxels. Here, we applied FSL’s cluster function to the statistical output from Randomise, whereby neighboring voxels for which the p-value exceeded our p<.01 threshold (across participants) were assigned to a cluster and the voxel-size of each cluster was summed. We then used AFNI’s 3dFWHMx method and the spatial AutoCorrelation function to conduct simulations to ask how often we would expect a cluster of a given voxel size based on chance alone. First, the smoothness of the original data on which the native searchlights were run was estimated, as was how correlated a given voxel was with its neighbors (i.e., -acf method). Second, the acf estimates were fed into the AFNI 3dClustSim function (Cox, 1996), which models the minimum number of contiguous voxels needed for a cluster to be deemed statistically significant at a voxel threshold of p<.01 and a cluster threshold of p<.05. In particular, 3dClustSim generated 10,000 artificial simulations (the default) with the same dimensions and smoothness as the searchlight dataset, but composed of noise derived from the residuals of the GLMs used to model the data inputs to the searchlight analysis (see above). The maximum cluster size was obtained for every random permutation. For the observed data, only cluster sizes that exceeded the 95^th^ percentile value of the random cluster sizes were considered significant (i.e., p<.05; two-sided, second-nearest neighbor).

## Results

### Children and adults learned overlapping pairs

Twenty-five adults (*M* = 19.06, *SD* = 3.36) and twenty-seven children between the ages of 7 and 10 years (*M* = 9.10, *SD* = 1.11) learned to associate individual objects with a person or place (Figure 1a).

The same person or place (six each) was paired with three different objects, for a total of 36 overlapping pairs. We used stimuli familiar to participants to encourage high-level performance (Methods), which in turn allowed us to ask about the neural organizational schemes that enabled successful memory for related events at different ages.

Moreover, before MRI scanning (Figure 1b), participants completed between two and five learning/retrieval repetitions until a 90% performance criterion was reached (Figure 1c; Supplementary Data).

Following learning, participants recalled the pairs during MRI scanning.

Participants were cued with an object followed by a 9-second delay, during which time they were instructed to hold the associated person or place in mind for an upcoming match/mismatch memory decision (Figure 1b). Both children and adults retrieved the overlapping memories, as evidenced by high and above-chance performance in both groups (all *t*s>25.39, all *ps*<.001; Figure 1d). Yet children still performed worse than adults (*t*(31.49) = 5.62, *p*<.001, d = 1.51, 95% CI = [0.06, 0.12]; Figure 1d). Thus, as a further step to isolating the neural organizational schemes and memory reinstatement mechanisms that enabled successful memory, unless otherwise noted, subsequent analyses were restricted to trials in which retrieval was successful during final learning and scanned recall (Methods; Supplementary Table 1 for trial counts).

### Children differentiate related memories

To assess organization of related memories in anatomical hippocampus, we used RSA (Kriegeskorte et al., 2008) to quantify the similarity among activation patterns evoked by object cues that were never directly observed together, but that shared an overlapping memory element (person/place; Figure 2a). In such cases, the related objects might evoke patterns that are significantly either more or less similar to one another as compared to unrelated objects. For such unrelated events, there are no shared links through which to organize the events, thus serving as an effective baseline.

**Figure 2.**
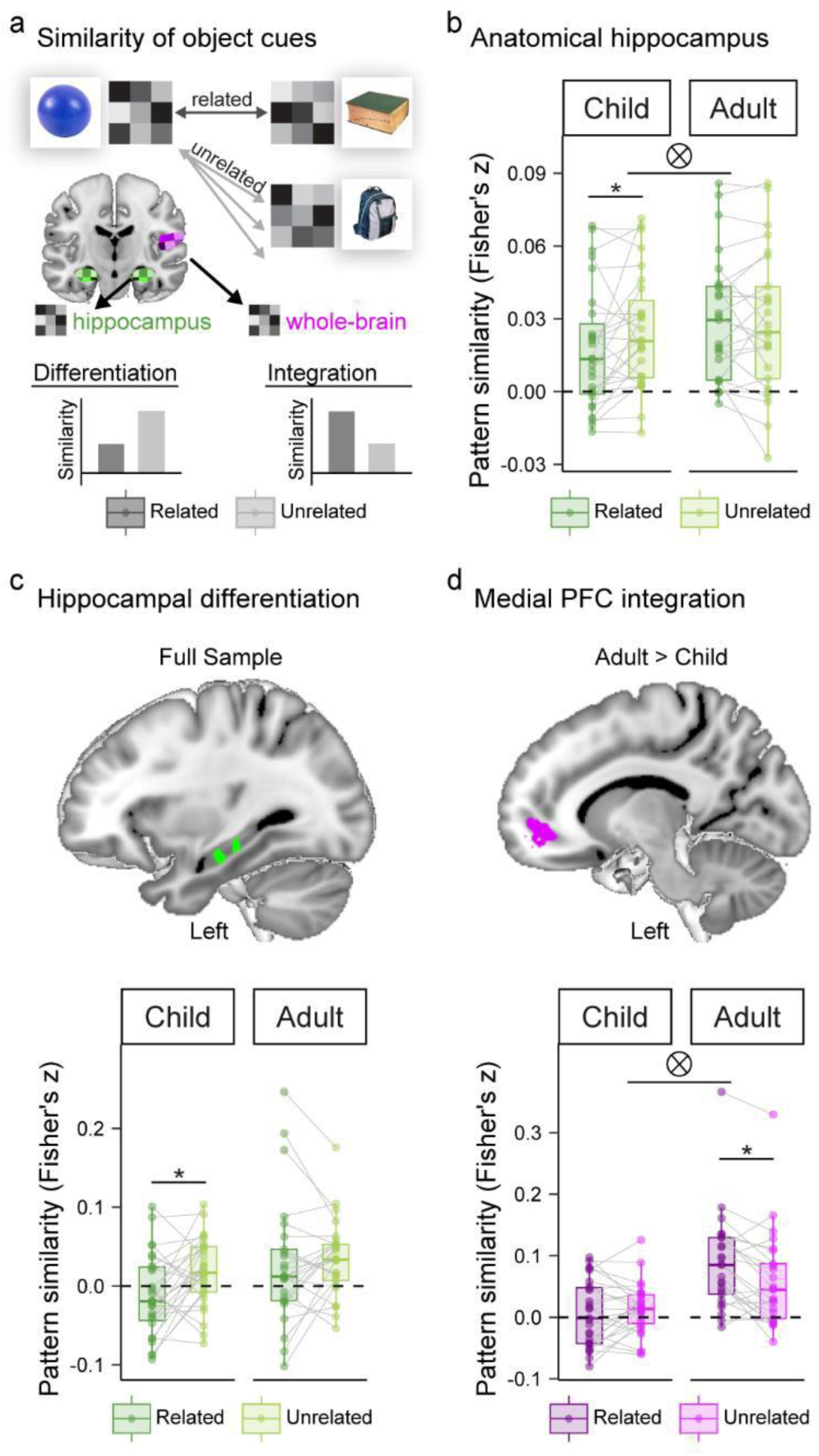
Children differentiate related object cues, while adults integrate. **a** Depiction of representational similarity analyses. We compared the similarity of fMRI patterns for related as compared with unrelated objects to assess evidence for differentiation (related < unrelated) and integration (related > unrelated). **b** Neural patterns extracted from bilateral anatomical hippocampus. Box plots reflect pairwise similarity values averaged across object cues that were related or unrelated, separately for participants within each age group. **c** Hippocampal searchlight regions showing a significant main effect of differentiation when examined across the full sample displayed on the 1-mm MNI template brain. Clusters are significant after small-volume correction for multiple comparisons within anatomical hippocampus. Interrogating effects within these clusters in each age group separately revealed significant differentiation in children but not adults (boxplot). **d** Whole-brain searchlight region showing a significant effect of integration in medial PFC which interacted with age group, with adults showing significantly greater integration than children. Cluster is significant after correction for multiple comparisons across the whole brain. Interrogating effects within this cluster in each age group separately showed that integration was significant in adults, but children showed no evidence of integration nor differentiation. This analysis also revealed a significant age interaction, in line with the fact that this searchlight cluster was defined based on this effect (marked here simply for clarity). Additional whole-brain results related to **c** and **d** are reported in Supplementary Table 2 and Supplementary Figures 2 & 3. For **b**-**d**, box plots depict the median (middle line), 25^th^ and 75^th^ percentiles (box), and the largest individual values no greater than the 5^th^ and 95^th^ percentiles (whiskers). Dots reflect individual participant means; lines connecting dots depict within-participant differences; n = 52 (27 children; 25 adults). Individual participant values (dots) that extend beyond the whiskers are considered outliers, defined as values that were 1.5 times greater than the interquartile range (IQR). Asterisks reflect a significant related/unrelated difference within an age group (\**p* < .05) while tensor product symbols indicate a significant interaction between age group and related/unrelated conditions at **b** *p* < .05 or **d** *p* < .001.

There was a trending difference in neural pattern similarity between related and unrelated objects (*F*(1,50) = 3.64, *p* = .06, *ηp*^2^ = .07) that was qualified by an interaction with age group (*F*(1,50) = 4.65, *p* = .04, *ηp*^2^ = .09), such that children (*F*(1,26) = 6.06, *p* = .02, *ηp*^2^ = .19, 95% CI = [-0.02, -0.002]) but not adults (*F*(1,24) = .05, *p* = .82, *ηp*^2^ = .002, 95% CI = [-0.005, 0.007]) exhibited hippocampal representations for related objects that were *less* similar than those for unrelated objects (Figure 2b). Notably, this differentiation pattern was absent in visual regions that may instead reflect perceptual similarities among objects (Supplementary Figure 1a) and in neocortical regions implicated in reinstatement of associated memory elements (Supplementary Figures 1b-c). Moreover, this effect remained even when we controlled for hippocampal BOLD activation during the cue (Supplementary Data). Our approach thus enabled us to uncover subtle differences in the organization of hippocampal memory traces not reflected in neocortex and that were above and beyond mean hippocampal activation.

The findings across our *a priori* anatomical hippocampal ROI establish that children, but not adults, differentiate related memories. However, we also know that the same memories can be represented as both differentiated and integrated within the same person, within different voxels across hippocampus and neocortex (Molitor et al., 2021; Schlichting et al., 2015), raising the possibility that different organizational schemes might be evident within different parts of hippocampus. In separate searchlight analyses, we therefore looked for voxels exhibiting differentiation or integration of related object cues in hippocampus and neocortex (Figure 2a). We searched for voxels that showed developmental differences in representational scheme (two-sample t-test: adult>child; child>adult), as well as voxels that showed a neural scheme across both groups (one-sample t-test of full sample vs. 0). To verify that significant coding effects were evidenced in both age groups separately for clusters identified across the full sample (>0), and for the group showing enhanced organization for clusters showing developmental differences (e.g., adults for adult>child contrast), for all significant clusters we additionally considered whether effects were present in each age group separately.

Results in hippocampus showed differentiation in children but not adults, but no evidence of integration. Specifically, we found no evidence of hippocampal integration in adults nor children when considered together; nor any clusters that showed developmental differences in either hippocampal integration or hippocampal differentiation. By contrast, clusters in left anterior hippocampus (Figure 2c; cluster center of gravity in the Montreal Neurological Institute (MNI) template coordinates (mm): *x*, *y*, *z* = -32.1, -18.7, -17.5) and left hippocampal body (Figure 2c: *x*, *y*, *z* = -29.7, -27.2, -12.8) showed significant differentiation across the whole group. However, it is noteworthy that when we interrogated these hippocampal clusters within the two age groups separately, we found that this effect was driven by significant or trend-level differentiation in the child group (anterior: *t*(26) = -2.27, *p* = .03, d = -0.44, 95% CI = [-0.076, -0.004]; body: *t*(26) = -1.99, *p* = .06, d = -0.38, 95% CI = [-0.080, 0.001]; average across clusters: *t*(26) = -2.90, *p* = .01, d = -0.56, 95% CI = [-0.068, -0.012]), with no evidence of differentiation just among adults (*p*s>.36) (Figure 2c; averaged across clusters). This pattern hints that while differentiation is the primary scheme for memory organization in children, it is not invoked in the adult group in the present task.

### Integration is more pronounced in adults

We next asked whether integration was more pronounced in the adult mPFC or elsewhere. Because different representational schemes have been previously observed in different subregions of mPFC (Schlichting et al., 2015), instead of using an anatomical ROI, here we chose a searchlight approach. In particulate, we conducted a whole-brain searchlight, asking whether mPFC or any other regions exhibited integration. As anticipated, the searchlight revealed that adults showed greater integration in mPFC (Figure 2d; two-sample t-test: adult>child; *x*, *y*, *z* = -15.30, 44.80, -4.43), as well as in superior parietal cortex and posterior cingulate (Supplementary Table 2; Supplementary Figure 2c). Within the regions showing integration in adults, children showed either no reliable/significant representational scheme (mPFC) or differentiation (parietal; posterior cingulate; Supplementary Figures 3c). In addition to revealing a general lack of evidence for integration in children (except for caudate; Supplementary Table 2 & Supplementary Figures 2b & 3b), this whole-brain searchlight further revealed that children (but not adults) showed differentiation in a number of other regions (Supplementary Table 2), with several of these clusters showing developmental differences, such that differentiation was enhanced in children relative to adults (Supplementary Figures 2 & 3). Together, these findings suggest that the neural substrates implicated in memory organization in adulthood may serve a different representational function earlier in life.

### Children and adults reinstate specific event features in neocortex

Having established that both children and adults systematically organized overlapping memories, with children relying on differentiation in hippocampus and neocortex and adults showing integration in neocortical regions including mPFC, our second goal was to test whether such discrepant schemes tracked neocortical reinstatement of the associated memory features. To address this question, we measured reinstatement at two levels of specificity: category (people vs. places) and item (Pinocchio vs. Peter Pan). We focused on functionally defined reinstatement regions of interest within anatomical VTC and parietal cortex (Figure 3a; Methods), guided by prior work (Fandakova et al., 2019; Trelle et al., 2020). Because we were only interested in effects wherein reinstatement was significantly greater than zero, all tests measuring above baseline reinstatement were one-tailed.

**Figure 3.**
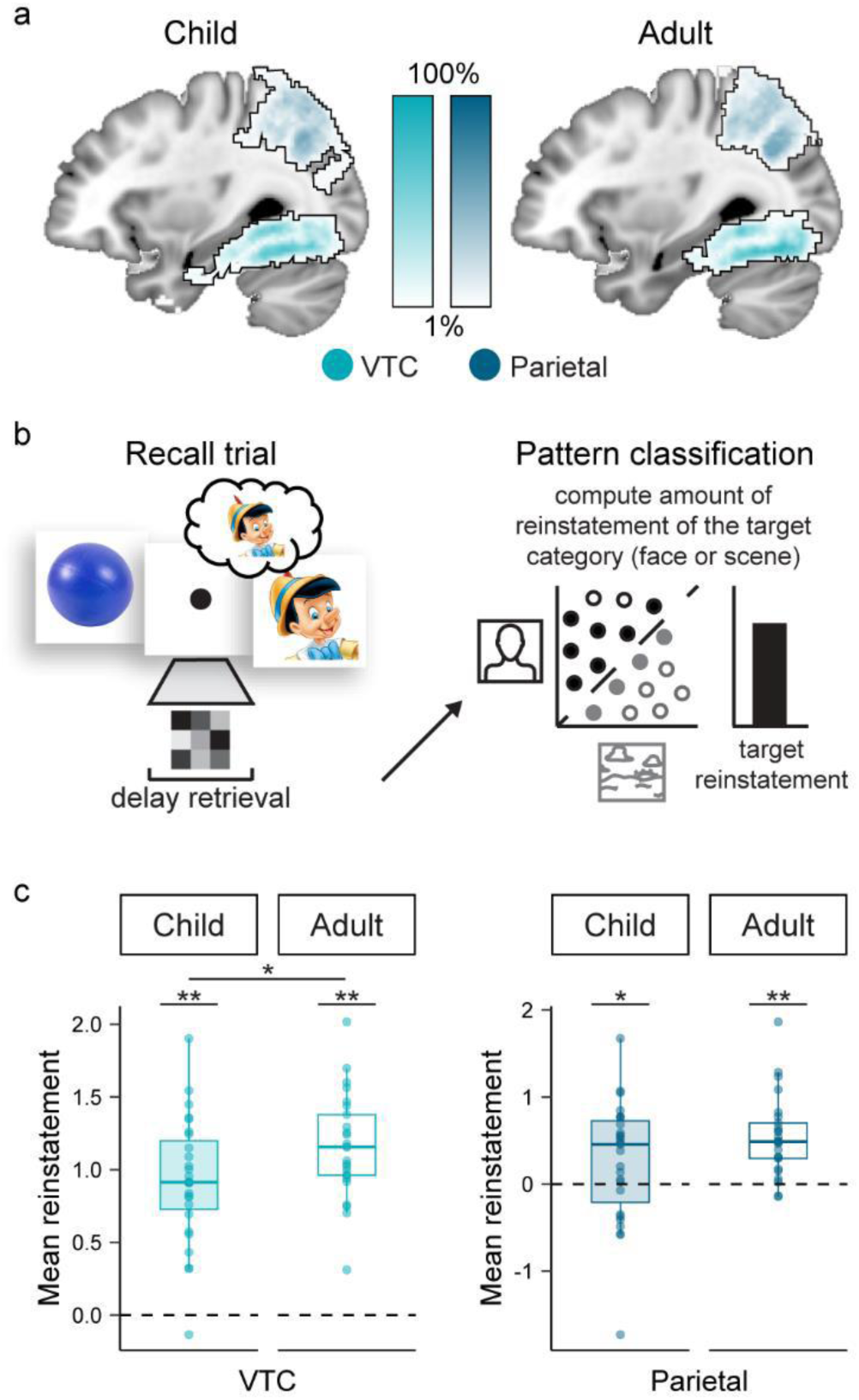
Children and adults reinstate the target category. **a** Voxels included within the VTC and parietal functional regions of interest (fROIs), separately for each age group. For each participant, we tested for reinstatement in the voxels that they engaged most during face and scene perception in a separate localizer task, within VTC and parietal cortex separately. Maps show the percentage of participants who had that same voxel in their fROI, plotted as a color-coded gradient. **b** Category decoding approach. We trained a multivoxel pattern analysis classifier to identify patterns of activity associated with face and scene processing in the VTC and parietal fROIs separately. For each delay retrieval pattern—which was modeled as a single voxelwise beta image across the 9-second retrieval period—we then applied the trained pattern-classification algorithm to estimate the amount of evidence for the target category (i.e., face or scene) reinstated. For each participant, we computed a single target reinstatement index by averaging the target classifier evidence (continuous probabilities) across trials, within each fROI separately. To correct for non-normality, the raw classifier output was transformed to logits. **c** VTC (left) and parietal cortex (right) category reinstatement. A reinstatement index reliably above 0 (i.e., the black dashed line) indicates significant target category evidence. Box plots depict the median (middle line), 25^th^ and 75^th^ percentiles (box), and the largest individual values no greater than the 5^th^ and 95^th^ percentiles (whiskers). Dots reflect individual participant means; n = 52 per plot (27 children; 25 adults), with dots extending beyond the whiskers reflecting outliers, defined as values that were 1.5 times greater than the interquartile range (IQR). Asterisks above each box reflect a significant within-group effect at a one-tailed threshold of p < .05 (*) or p < .001 (**), while asterisks between bars reflect a significant between-group effect at a two-tailed threshold of p < .05.

To quantify the degree to which the target category was reinstated during the delay period, we trained an MVPA classifier (Norman et al., 2006) to identify patterns of activity associated with face and scene processing within each functionally-defined reinstatement ROI (Methods; Supplementary Data: Classifier validation accuracy; for consideration of an RSA-based category measure instead, see Supplementary Figure 4). We then applied the trained classifier to patterns of fMRI activation measured during the 9-second delay interval (Figure 3b), to quantify the amount of face and scene evidence (Methods). Both age groups showed categorical reinstatement in VTC (adult: *t*(24) = 15.94, p < .001, d = 3.19, 95% CI = [1.01, 1.30]; child: *t*(26) = 11.24, p < .001, d = 2.16, 95% CI = [0.76, 1.09]; one-tailed) which further differed by age (*t*(50) = 2.08, p = .04, d = 0.58, 95% CI = [0.008, 0.452]; two-tailed), such that adults showed higher-fidelity reinstatement (Figure 3c). Adults and children additionally showed reinstatement in parietal cortex (adult: *t*(24) = 5.82, p < .001, d = 1.17, 95% CI = [0.35, 0.73]; child: *t*(26) = 1.95, *p* = .03, d = 0.38, 95% CI = [-0.01, 0.53]; one-tailed; Figure 3c), which did not differ by age (*t*(50) = 1.71, p = .09, d = 0.48, 95% CI = [-0.05, 0.61]; two-tailed).

We next examined item-level reinstatement. Specifically, we used RSA to quantify the similarity between patterns of fMRI activation measured during retrieval of a target item and viewing of the same item during a separate pre-exposure task (same-item similarity; Figure 4a), as compared to viewing other items from the same visual category (different-item similarity; Methods). Item-level reinstatement in VTC was significant in adults (*t*(24) = 2.32, *p* = .01, d = 0.47, 95% CI = [0.001, 0.011]; one-tailed) but only a trend in children (*t*(26) = 1.52, *p* = .07, d = 0.29, 95% CI= [-0.001, 0.009]; one-tailed), and was not significant at either age in parietal cortex (*t*s < 1.13; *p*s > .14; one-tailed). Neither VTC nor parietal item reinstatement differed as a function of age group (*t*s < 0.52; *p*s > .61). Given only modest item reinstatement effects in the category-selective reinstatement ROIs (being observed only in VTC and only for adults), we additionally conducted a whole-brain item-level reinstatement searchlight analysis, which was small-volume corrected within the VTC and parietal reinstatement ROIs as well as within hippocampus given prior adult work (Mack & Preston, 2016). For both groups, we found significant item reinstatement in more focal regions within the functionally-defined reinstatement ROIs as well as elsewhere in the brain (Supplementary Table 3; Supplementary Figure 5), though developmental differences were observed in hippocampus, with the locus of item reinstatement shifting from posterior to anterior between childhood and adulthood (Supplementary Figure 6).

**Figure 4.**
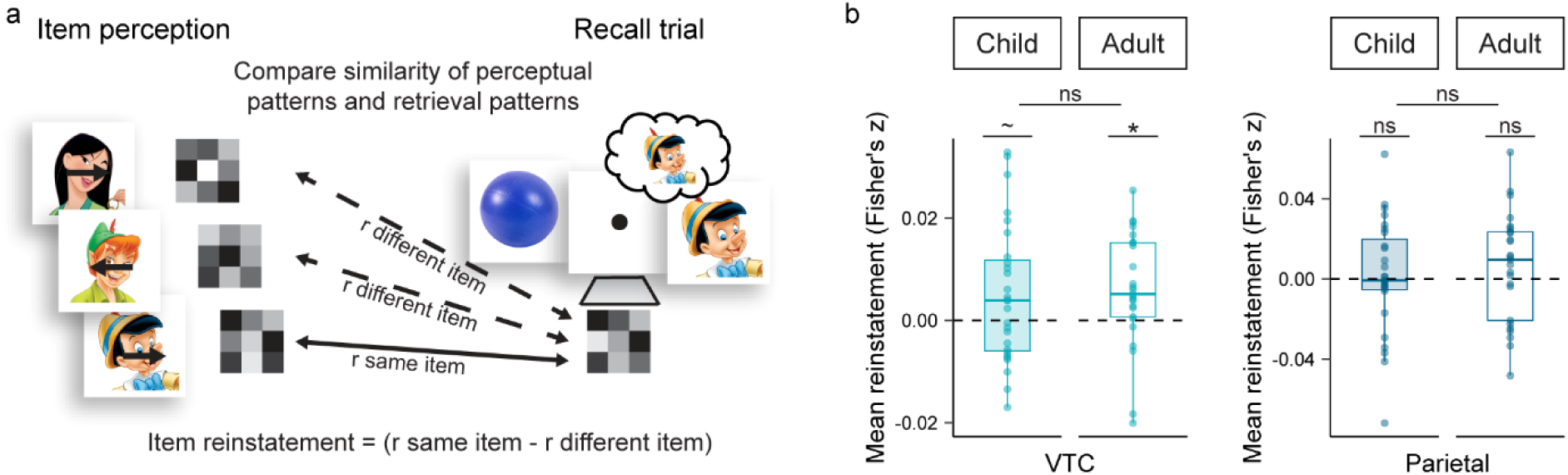
Children and adults reinstate target items. **a** Representational similarity analysis approach. Representational similarity analysis was implemented at the level of the full category-selective functional regions of interest (fROIS; from Figure 3a) as well as at the individual voxel level via separate searchlight analyses (see Supplementary Table 3 & Supplementary Figures 5 & 6). During item perception/the pre-exposure task, participants indicated whether an arrow superimposed on each stimulus (face/place) was pointing left or right. For each scanned recall trial, we correlated fMRI activation patterns from the delay period with viewing of the same item during a pre-exposure item perception localizer (same-item; solid arrow), which was compared to viewing of different items from the same visual category (different-item baseline; dashed arrow). For each participant, we computed a single item reinstatement index by averaging the mean same item – different item similarity difference across recall trials. **b** Pattern similarity results within VTC (left) and parietal cortex (right) fROIs. A mean reinstatement index reliably above 0 indicates significant item evidence, reflecting greater same-item similarity as compared to the different-item baseline. Box plots depict the median (middle line), 25^th^ and 75^th^ percentiles (box), and the largest individual values no greater than the 5^th^ and 95^th^ percentiles (whiskers). Dots reflect individual participant means; n = 52 per plot (27 children; 25 adults), with dots extending beyond the whiskers reflecting outliers, defined as values that were 1.5 times greater than the interquartile range (IQR). Symbols above each box reflect a significant within-group effect at a one-tailed threshold of p < .05 (*), or that is at trend levels (∼). The letters “ns” above and between bars indicate a nonsignificant within-group or between-group effect, respectively.

We also wanted to confirm that modest evidence of item reinstatement in the VTC and parietal ROIs was not an artifact of the long delay, particularly in children. To address this possibility, we performed additional analyses in which we separately considered reinstatement during the first and second halves of the delay period in both age groups (Methods). There were two possible ways in which reinstatement might differ by half. The first possibility was a decrease in reinstatement across the delay, suggesting that the delay may be too long for children to hold the reinstated memory in mind. The second possibility was that we would instead see an increase in reinstatement, which would support the idea that it took children time to reinstate the memory (as we anticipated [Hajcak & Dennis, 2009]; Methods). Our results of this supplementary analysis were generally more consistent with the latter of the two possibilities, showing that both children and adults exhibited evidence of reinstatement across the full 9-second delay, with category (and some item) reinstatement evidence being stronger in the second half of the delay immediately prior to the decision probe (Supplementary Figure 7).

### High-fidelity neocortical reinstatement is linked to mnemonic decision making

One important clue as to whether neocortical reinstatement directly guides memory behavior comes from asking how it relates to decision behavior on a trial-by-trial basis, in which participants were asked to judge whether the retrieved memory matched the decision probe (Figure 1b). To test this, we conducted a mixed effects and drift diffusion models to examine whether variability in item (Figure 4b, same-item similarity) and category (Figure 3c) reinstatement during the delay predicted the probability of making a correct and/or faster response within a participant on a given trial (Methods). Because our mixed effects model tested which reinstatement signatures (VTC item, VTC category, Parietal item, Parietal category; in a single model) were associated with correct versus incorrect performance, we did not restrict our analyses to correct trials.

We found that more strongly reinstated memories were associated with more accurate responses (VTC category reinstatement: χ^2^1 = 4.12, *p* = .04, Figure 5a, left; trend for parietal cortex item reinstatement: χ^2^1 = 3.52, *p* = .06, Figure 5a, right), which were significant predictors of performance despite the fact that our model additionally accounted for the main effect of age (χ^2^1 = 29.08, *p* < .001; model comparison: AICbase= 848.16, AICMainEffect= 817.74, χ^2^1 = 32.42, *p* < .001; a more complex model that allowed reinstatement to interact with age did not significantly improve model fit, χ^2^1 = 0, *p* = 1.00), as well as the potential influence of univariate activation, including cue-period hippocampal BOLD activation (χ^2^1 = 3.47, *p* = .06) and delay-period VTC and parietal BOLD activation (χ^2^1 < 0.24, *ps* > .63). Follow-up analyses of each age group separately showed that this effect was found in children (VTC: χ^2^1 = 3.62, *p* = .06; Parietal: χ^2^1 = 3.42, *p* = .06) but not adults (ps > .56).

**Figure 5.**
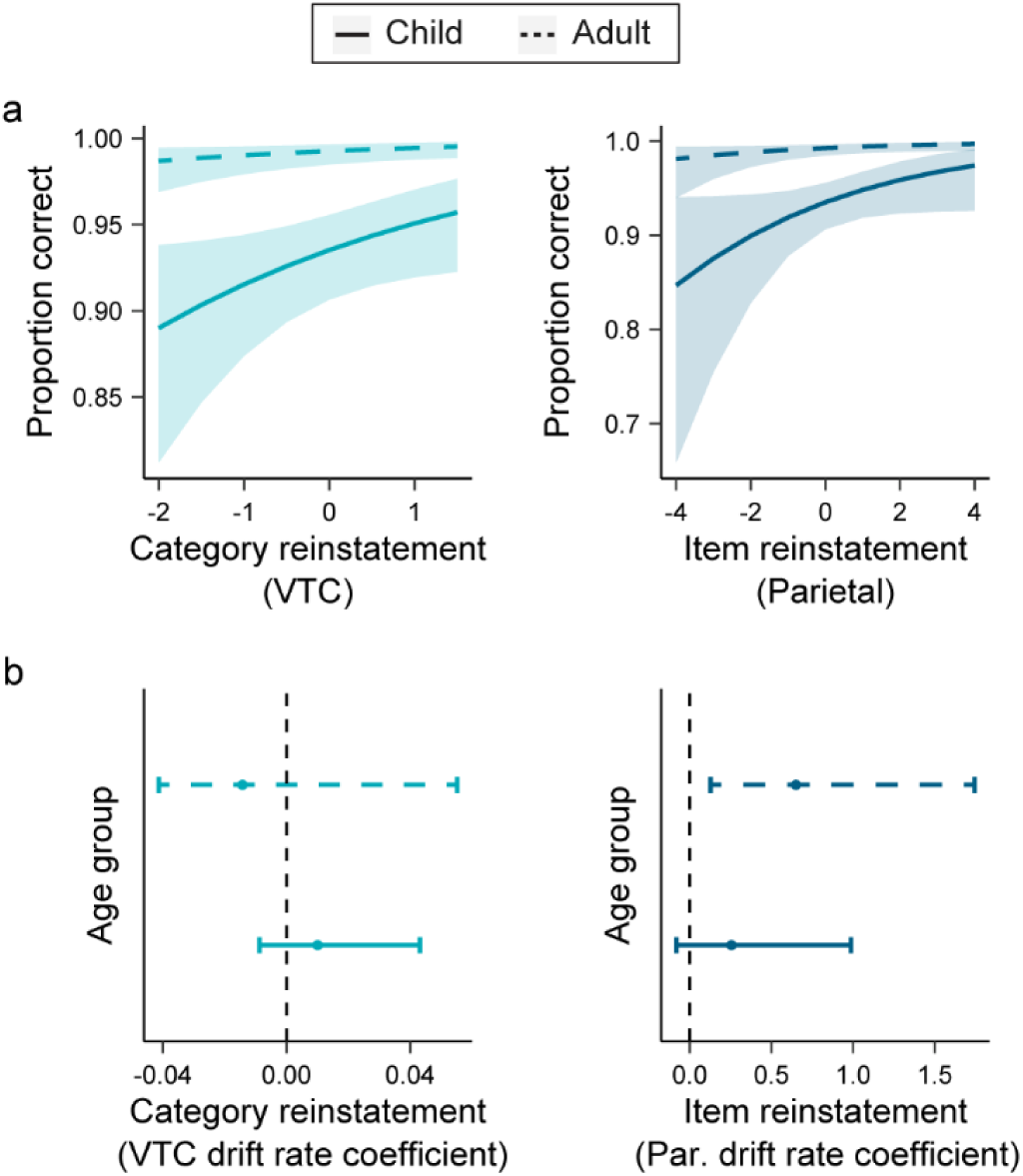
Enhanced neocortical reinstatement in children and adults promotes superior memory behavior. We tested the within-participant association between raw same-category and same-item reinstatement during the delay period with the probability of making either a correct **a** or speeded **b** match/mismatch behavioral response. **a** Results of mixed effects model examining decision accuracy. Here, significant predictors are depicted in separate plots for visualization purposes only, as all predictors (see Method) were included in a single statistical model. All predictors were scaled and centered within participants to remove participant-specific effects. For both plots, n = 52 (27 children; 25 adults); age group was modeled as a main effect. We found that greater category-level reinstatement in VTC (left) and a trend for greater item-level reinstatement in parietal cortex (right) each predicted successful performance. Lines (solid and dashed) reflect model predictions, while shaded areas reflect 95% confidence interval around the model predictions. **b** Results of drift diffusion models (DDM) examining decision speed. Here, the VTC category and parietal cortex item reinstatement indices were entered into a single DDM model of response speed, modeled as a linear effect on the drift rate parameter that reflects the rate of evidence accumulation (with two separate models run, one each for children and adults). Whereas higher drift rate coefficients reflect faster evidence accumulation and response time; lower values reflect slower evidence accumulation and response times. The interaction of drift rate and reinstatement in VTC (left) and parietal cortex (right) are depicted as forest plots. The lines represent 95% confidence intervals of the 2,000 posterior parameter estimates (one-tailed), with circles representing the mean of the posterior parameter distributions. Greater item reinstatement in parietal cortex predicted faster response times, but only in adults, which was reflected by drift rate coefficient confidence intervals not encompassing and greater than 0. No significant effects were observed for category reinstatement in VTC, in either adults or children.

Likewise, in adults (who reached ceiling levels of accuracy) but not children, faster responses were associated with more strongly reinstated memories (Figure 5b, right; parietal item) while controlling for hippocampal BOLD activation during the cue which was shown to be a significant predictor of accuracy above. Furthermore, in line with the observation that reinstatement increased over the delay, reinstatement at the end (but not beginning) of the delay was behaviorally significant, predicting response accuracy in children and speed in adults (Supplementary Figure 8).

### Different representational schemes have benefits for reinstatement at different ages

We found that children and adults deploy high-fidelity neocortical representations (Figures 3 & 4) in service of mnemonic decisions (Figure 5). However, whereas children rely on hippocampal differentiation to organize related memories (Figure 2b), adults relied on integration, including in mPFC (Figure 2d) as anticipated. In a final set of analyses, we thus tested whether the degree of hippocampal differentiation and mPFC integration was associated with neocortical reinstatement in children and adults, respectively. In addition to our *a priori* hypothesis regarding mPFC integration, we further explored the relationship between neocortical reinstatement and other searchlight regions that revealed significant integration, either in adults (parietal cortex, posterior cingulate) or children (caudate). When quantifying delay-period reinstatement, we focused on only those neocortical functionally-defined reinstatement ROIs that, in both age groups and across the full delay, either showed (1) reliable reinstatement (i.e., category in VTC and parietal cortex; Figure 3c) or for which (2) trial-wise variability in reinstatement was linked to behavior (i.e., item reinstatement in parietal; Figure 5) at significant or trend levels.

Memory organization in hippocampus (anatomical) and mPFC (searchlight cluster) were significantly associated with category-level reinstatement in VTC for children and adults, respectively. Specifically, hippocampal differentiation was associated with enhanced reinstatement in children (*t*(57.15) = -2.29, *p* = .03) but not adults (*t*(43.59) = 0.43, *p* = .67; relationship differed significantly between the two groups [Interaction: χ^2^1 = 3.78, *p* = .05; AICbase=10042, AICinteraction=10041, χ^2^2=4.75, *p*=.09; Figure 6a]). Conversely, mPFC integration was associated with enhanced reinstatement in adults (*t*(43.31) = 2.36, *p* = .02) but not children (*t*(57.84) = -1.32, *p* = .19; Interaction: χ^2^1 = 6.69, *p* = .01; AICbase=10043, AICinteraction=10042, χ^2^3=7.52, *p*=.06; Figure 6b). In addition, mPFC integration was associated with item-level reinstatement in parietal cortex (χ^2^1 = 7.77, *p* = 0.005; best-fitting model was one that did not include age group [i.e., base model; Main Effect and Interaction model fits: ps > 0.28]); this same relationship was not evidenced for hippocampal cue organization (χ^2^1 = 2.39, *p* = 0.12; base model). Further, neither hippocampal nor mPFC cue organization was associated with category reinstatement in parietal cortex (hippocampus: χ^2^1 = 2.02, *p* = 0.16; mPFC: χ^2^1 = 0.93, *p* = 0.33; both base models).

**Figure 6.**
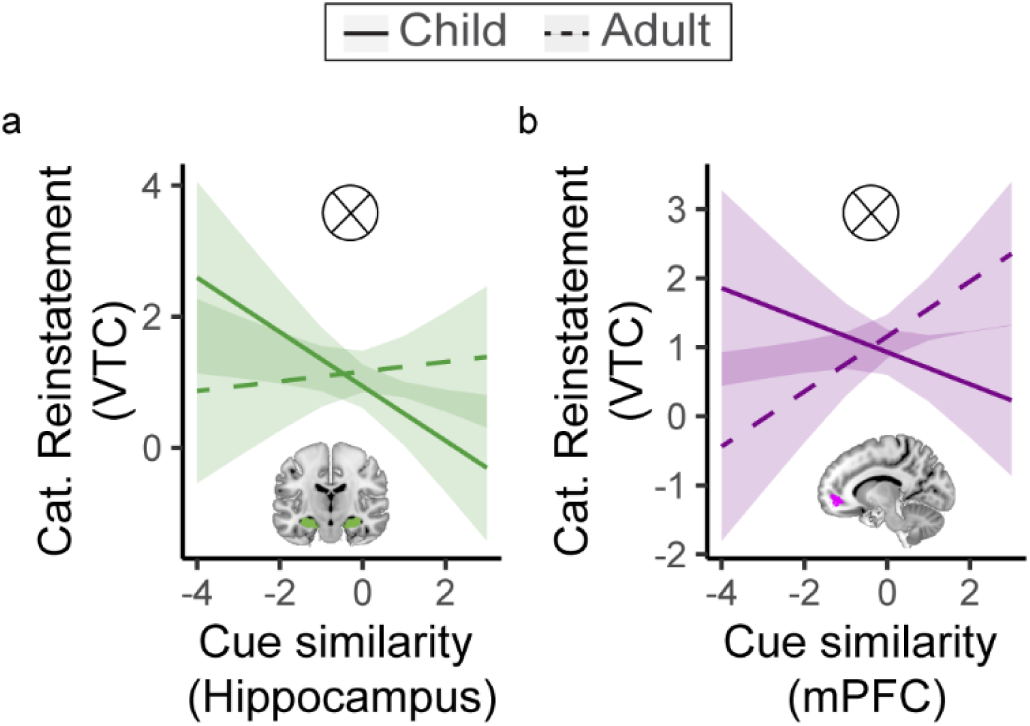
The hippocampal and mPFC representations that governed how related memories were organized tracked the degree of ensuing neocortical reinstatement of the associated face and scene elements at the within-person level. In both panels, for each participant and retrieval trial separately, we computed the averaged similarity of a retrieval cue object with related cue objects that shared the same face or scene associate (Figure 2a; related similarity). Thus, whereas lower cue similarity values reflect greater dissimilarity of related memories, higher values reflect greater similarity. We tested the association between that cue similarity value with reinstatement during the corresponding delay period for that same retrieval trial (Figure 3b; Target reinstatement example trial). **a** In children (solid green line) but not adults (dashed green line) there was a negative trial-wise association between cue similarity in anatomically-defined bilateral hippocampus (Figure 2b) and category reinstatement in VTC. In other words, lower similarity in hippocampus was associated with higher-fidelity reinstatement of the target category. **b** In adults (dashed purple line) but not children (solid purple line) there was a positive trial-wise association between cue similarity in the mPFC region defined from the searchlight (Figure 2d) and category reinstatement in VTC. That is, greater similarity in mPFC was associated with higher-fidelity reinstatement of the target category. For both panels (each comprising a different model), age group was modeled as an interaction and both interaction terms were significant at a threshold of *p* ≤.05 (marked here with a tensor product symbol). For each model, predictors (x-axis) were scaled and centered within participants to remove participant-specific effects. Lines (solid and dashed) reflect model predictions, while shaded areas reflect 95% interval around the model predictions.

Finally, in contrast to mPFC which revealed a significant association between memory integration and VTC category reinstatement among adults, there was no such association between cue organization and VTC category reinstatement in other regions that showed significant memory integration (Supplementary Figures 2 & 3), including in parietal cortex (Interaction: χ^2^1 = 3.09, *p* = .08, Adult slope: *p* = 0.12, Child slope: *p* = 0.35; Interaction model fit: p = .13), posterior cingulate (Interaction: χ^2^1 = 5.06, *p* = .02, Adult slope: *p* = 0.07, Child slope: *p* = 0.19; Interaction model fit: p = .05), or caudate (Base model: χ^2^1 = 0.37, *p* = .54; Main Effect and Interaction model fits: ps > 0.31).Together, the findings suggest that different hippocampal and mPFC organizational schemes are uniquely and differentially beneficial for reinstatement at different ages.

## Discussion

Prominent developmental theories have long proposed that age-related improvements in learning and cognition are rooted in the emergence of an increasingly complex representational system (Karmiloff-Smith, 1986). Yet despite both its theoretical importance (Bauer & Varga, 2017; Keresztes et al., 2018) and educational relevance (Varga et al., 2019), the question of how children—for whom the neural substrates supporting memory are still developing (Bauer et al., 2019; Brod et al., 2017; Fandakova et al., 2017; Ghetti & Bunge, 2012; Keresztes et al., 2018; Ofen et al., 2007; Østby et al., 2009; Schlichting et al., 2017; 2022; Simmonds et al., 2014)—represent related events to enable later memory retrieval remains unanswered. By leveraging fMRI with representational analysis methods, we show that children code related memories via hippocampal differentiation, wherein events that share overlapping features are represented as *less* similar to one another compared to unrelated events. In contrast, adults deploy an alternate representational scheme in which memories for related experiences are integrated, including within mPFC, wherein related events are more similar to one another than they are to unrelated events. Critically, these representational schemes were associated with memory reinstatement in children and adults, respectively: That is, hippocampal differentiation in children and mPFC integration in adults each tracked higher-fidelity neocortical reinstatement, which in turn was linked to superior memory behavior in both groups. Together, these findings indicate that children may be biased toward disambiguating overlapping experiences, while adults build representations that code common features across them. Such a shift in memory organization may confer certain behavioral advantages during each period in development.

Our primary objective in designing the present study was to isolate how the developing brain organizes overlapping experiences in ways that enable successful, high-fidelity retrieval of individual memories. Although children often perform less accurately than adults when both disambiguating between (Ngo et al., 2018; Rollins & Cloude, 2018) and integrating across (Bauer et al., 2021; Schlichting et al., 2017; Shing et al., 2019; Wilson & Bauer, 2021) overlapping events, such developmental differences need not reflect a fundamental inability to retrieve overlapping experiences. Indeed, children as young as 6 years can successfully recall and connect related experiences (Bauer & San Souci, 2010), while discrimination of highly similar memories becomes adult-like by approximately 10 years (Rollins & Cloude, 2018). These behavioral findings hint at important changes in the neurobiological system that supports representation of related events and highlights middle childhood (7-10 years) as a candidate window in which the memory system has achieved at least some adult-like functionality. Yet studies of behavior cannot tell us how children successfully represent and recall related experiences, or whether they do so based on the same or different types of memory representation as evidenced in adults. By quantifying representational similarity of related events as they were recalled from memory, the present study thus offers a novel window into the representational competencies available within the early hippocampal and mPFC memory systems.

We found that hippocampal differentiation, but not integration, was associated with successful learning and retrieval of related memories in children. Our finding joins one prior study showing hippocampal differentiation in children of a similar age (Benear et al., 2022). One notable extension of the prior work is that here we measured neural representations only following multiple learning and retrieval attempts, which enabled us to identify the neural organizational schemes that underpinned successful memory behavior. In contrast, the neural representations in prior research (Benear et al., 2022) were assessed during initial exposure, and moreover, subsequent memory for the related experiences was not assessed. As the prior study showed children exhibited hippocampal differentiation only for previously experienced stimuli (i.e., familiar movies; Benear et al., 2022), but not for newly encoded movies, we suggest that here the multiple learning and test repetitions may have encouraged robust differentiation in children. Indeed, we know that children struggle to spontaneously organize related memories during learning (Abolghasem et al., 2023; Bauer et al., 2015; Varga & Bauer, 2013). Thus, by interleaving learning exposures with explicit retrieval tests, thereby strengthening overlapping memory formation, our study provides key insight into the endpoint of learning. In doing so, our results reveal hippocampal differentiation—an active coding strategy that might help resolve interference among related memories (Chanales et al., 2017; Favila et al., 2016; Hulbert & Norman, 2015)—as a means through which the still-developing brain organizes experience, and ultimately, achieves successful neocortical reinstatement and retrieval behavior. The present results thus reveal new insights into the relationship between memory representation and behavior in children that could have implications for our understanding of how to promote overlapping memory, key predictors of educational success (Varga et al., 2019).

Beyond hippocampus, children in our study showed differentiation broadly across many neocortical regions. Such a tendency towards differentiation—or in some regions, an entire lack of systematic organization, perhaps akin to pattern separation, wherein non-overlapping traces arise through automatic orthogonalization (Norman & O’Reilly, 2003)—contrasted with the representations we observed in adults, which were largely integrated. Importantly, the observed lack of neural integration in children aligns with past evidence (Bauer et al., 2021; Schlichting et al., 2022; Wilson & Bauer, 2021) and cannot be explained by age-related differences in either behavior (Sastre et al., 2016) or prior experience (Benear et al., 2022), as has been suggested to drive other age-related activation and representational differences reported in the literature. Indeed, one exception to this overwhelming bias in children toward differentiation was in caudate, which showed integration. Importantly, such evidence helps to constrain our interpretation of null integration effects elsewhere, suggesting that, should children have formed this type of representation in either hippocampus or mPFC, our neural analytic approach was sufficiently powered to detect it.

Such an observation of integration in caudate underscores that children are not fundamentally incapable of this coding scheme, but rather that it may simply be less prevalent in their brains, primarily appearing in regions that mature earlier than neocortex (Van den Bos, Cohen, Kahnt & Crone, 2012). We speculate that the isolation of integration to caudate could indicate a role for simple action-outcome learning during the feedback phase of the three-alternative forced choice test (Van den Bos et al., 2012). As a reminder, to assess initial learning of the individual pairs, participants were cued with an object and instructed to choose the face or scene that was paired with it. By providing corrective feedback after each choice (action), we may have drawn children’s attention to the shared face/scene “outcome” (which appeared in the center of the screen) across test trials. Caudate aside, our finding of overwhelming neocortical differentiation might suggest that children’s neural biases can be characterized as shifting from differentiated (or pattern separated) to integrated over developmental time.

The observed age-related shift from differentiation toward integration is consistent with recent behavioral studies indicating pronounced developmental differences in reasoning behaviors that rely on extracting novel relations among memories. Although children are able to combine related memories to derive new factual knowledge (Bauer & San Souci, 2010) or make inferential reasoning judgments (Schlichting et al., 2017)—behaviors that may seem to imply and indeed (in adults [Molitor et al., 2021; Schlichting & Preston, 2014; Varga & Manns, 2021; Zeithamova et al., 2012]) benefit from neural integration—children may be accomplishing these same tasks using fundamentally different types of memories relative to adults (Bauer et al., 2021; Varga et al., 2019; Wilson & Bauer, 2021). For example, in contrast to adults, children often do not connect related facts or events during learning either spontaneously (Bauer et al., 2021; Wilson & Bauer, 2021) or even when instructed to do so (Abolghasem et al., 2023; Bauer et al., 2015; Schlichting et al., 2022). By contrast, similar instructions provided immediately before an inference test do facilitate performance (Bauer et al., 2015), suggesting an instruction facilitates interrogation of separate memories during flexible decision making.

Such behavioral data coupled with the differentiated neocortical organization observed here therefore points to the possibility that performance on reasoning tasks may be achieved by different underlying neural memory representations in children and adults (Kumaran & McClelland, 2012). Specifically, whereas adults may access an integrated representation directly to facilitate behavior, children may instead retrieve individual (differentiated or separated) representations and later recombine them when explicitly prompted to make an inference judgment (Bauer et al., 2017; Bauer & Varga, 2017; Schlichting et al., 2022; Wilson & Bauer, 2021). As we did not directly assess participants’ inferential reasoning or knowledge extraction behaviors, how such memory flexibility may be supported by different representational schemes across development remains to be seen. However, here we do see that differentiation improved reinstatement of directly learned events in children, whereas integration was beneficial in adults, thus revealing that different organizational schemes are beneficial at distinct ages.

In contrast to children, adults showed evidence for neural memory integration in several neocortical regions including mPFC. Importantly, such neural integration emerged despite no explicit demand to link related associations in memory. Indeed here, participants were instructed that they would need only to remember individual pairs, and in fact there was no behavioral requirement that participants make the novel connections (as in e.g., inferential reasoning). Such evidence of neural integration is consistent with both computational models of inference (Ritvo, Nguyen, Turk-Browne & Norman, 2023) and extant empirical data from adults (Molitor et al., 2021; Schlichting et al., 2015). For example, when trained to perform a task very similar to the one employed here, neural network models learn indirect connections between objects that share features and outcomes without reinforcement (Morton et al., 2020). In other words, integration is an emergent property of these models and parallels neural organization in human mPFC. Likewise, in prior adult work, integration has been observed during/after overlapping pair learning but before any explicit demand to make links among memories in a behavioral inference task (or even awareness that such a test is forthcoming; Molitor et al., 2021; Schlichting et al., 2015; Schlichting & Preston, 2014; Varga & Manns, 2021; Zeithamova et al., 2012). Interpreting the present data in the context of this prior literature, we thus conclude that a mature mPFC not only represents directly experienced associations, but further encodes the inferred latent structure of a task or an individual’s environment (Chan et al., 2016). Certain other aspects of our study, such as the use of well-known stimuli, may have further encouraged mPFC integration as participants could incorporate the new associations into existing memory networks (Bein et al., 2020).

In contrast to the mPFC integration findings, adults did not show significant evidence of either integration or differentiation in hippocampus. While this null effect may on its surface appear to conflict with prior adult studies (Molitor et al., 2021; Schlichting et al., 2015), we suggest that it does not; rather, these findings can be reconciled by considering the nuanced set of individual differences and situational factors (Zeithamova & Preston, 2017) that promote integration or differentiation in the mature brain. With respect to individual differences, even when holding the learning scenario constant, adults appear to vary in their representational tendencies (Schlichting & Preston, 2014; Varga & Manns, 2021; Zeithamova et al., 2012). For instance, white matter integrity between hippocampus and mPFC predicts individuals’ ability to link related events in memory (Schlichting & Preston, 2016). Along with other stable, individual differences in brain structure (Bauer et al., 2019; Schlichting et al., 2015; 2017), differences in strategy use (Richter et al., 2016), working memory capacity (Varga et al., 2019), and even culture (Millar, Serbun, Vadalia, & Gutchess, 2013) may change how individuals organize related memories. In the present study, significant variability across participants for any of these factors would preclude observing reliable integration or differentiation across our group of adults on average, without undermining the interpretation of our developmental findings.

An additional and perhaps related possibility for the null hippocampal findings in adults relates to the fact that the mature hippocampus is flexible and can form either differentiated or integrated representations, sometimes in parallel across different subregions (Molitor et al., 2021), depending on the specifics of the learning situation.

For example, one study (Schlichting et al., 2015) showed (anterior) hippocampal integration when initial associations were learned to a high level of accuracy before overlapping pairs were introduced, yet (both anterior and posterior) differentiation when overlapping associations were instead studied in an interleaved order. Notably, in that study, distinct regions of mPFC exhibited integration regardless of learning schedule, underscoring that it may complement more flexible hippocampal codes, with mPFC being comparatively less sensitive to task features. Thus, one possibility is that the interleaved training and more complex associative structure (i.e., three overlapping pairs as opposed to two) biased the adult hippocampus but not mPFC away from integration, while at the same time providing insufficient exposures to yield strong and consistent hippocampal differentiation (Chanales et al., 2017; Favila et al., 2016).

Two other neocortical regions—parietal cortex and posterior cingulate—showed memory organization schemes that differed significantly across age group, with children showing differentiation and adults integration. In other words, the same voxels in these regions served diametrically opposite representational functions across development.

Such findings generally align with past work. One possibility suggested by recent adult MRI data (Baldassano et al., 2018; Morton et al., 2020; Pudhiyidath et al., 2022) is that both parietal cortex and posterior cingulate play privileged roles in forming and utilizing relational learning structures. For example, posterior cingulate in adults has been implicated in forming knowledge structures that reflect spatiotemporal regularities (Baldassano et al., 2018; Pudhiyidath et al., 2022) while parietal cortex enables “shortcuts” through representational space to support rapid memory-guided decisions (Morton et al., 2020). At the same time, others have speculated that parietal cortex is an episodic memory buffer that accumulates mnemonic evidence for decision-making (Vilberg & Rugg, 2008; Zhang, Yin & Yang, 2022). In the present study, parietal cortex may index accumulation of the differentiated representations in children (from hippocampus) and the integrated representations in adults (from mPFC), prior to reflecting accumulation of the associated face/scene during the delay. However, note that here, in contrast to hippocampus and mPFC, parietal organization during the cue did not predict reinstatement of the associated face or scene during the ensuing delay, which might be expected under such an explanation. Interestingly, very few studies have interrogated parietal or posterior cingulate development, and those that did tended to focus on the development of metamemory processes (Fandakova et al., 2017; Tamnes et al., 2013). The present findings thus build on this work, suggesting that children and adults may rely on similar neocortical sites to activate a relevant memory space, the nature of which fundamentally differs as a function of age-related capacities in hippocampus and mPFC.

Our neural pattern analysis approach allowed us to detect memory reinstatement in both children and adults, which was moreover linked to behavior in both groups. Specifically, children and adults alike showed evidence of category- and item-level reinstatement of associative memory elements in neocortex. Notably, whereas adults showed broad evidence of item reinstatement in the VTC ROI as a whole across the entire delay, item-level reinstatement in children was observed at significant or trend-levels (1) across the whole VTC ROI, but only during the second half of the delay (Supplementary Figure 7; trend) or (2) in a number of searchlight clusters across the entire delay (Supplementary Figures 5 & 6). Beyond neocortex, we further showed that both children and adults reinstate specific item memories in hippocampus. Consistent with the view that that memory processes are differently distributed in the immature hippocampus (DeMaster & Ghetti, 2013), we showed that children recruited only posterior hippocampus, in contrast to adults who recruited anterior hippocampus (Supplementary Figure 6). Together these findings suggest that it is possible to quantify the internal contents of specific memories in children, provided the methods are sensitive to its relatively more circumscribed physical location and more delayed temporal profile. While prior work has shown that children and adults form similar neocortical representations of specific items at encoding (Fandakova et al., 2019), here we extend this finding to show comparable representations at retrieval. Moreover, the degree to which the associated faces/scenes were reinstated in neocortex, particularly during the second half of the delay, had relevance for memory-based decision making. More specifically, greater category-level VTC and item-level parietal reinstatement predicted successful decisions in children, whereas greater item-level parietal reinstatement predicted faster decision speed in adults. As discussed above, these findings are generally consistent with the view that parietal cortex plays a privileged role in accumulating mnemonic evidence in service of decision-making (Zhang et al., 2022).

Theories of memory development have long speculated that memory in childhood depends on the integrity of the hippocampal-neocortical system (Ghetti & Bunge, 2012), and the strength with which distributed memory traces are bound together, in particular (Howe & O’Sullivan, 1997). Therefore, having established the behavioral relevance of neocortical reinstatement, our final question was whether the hippocampal and mPFC representational schemes employed to organize overlapping memories tracked the fidelity with which those associated memory elements were reinstated in neocortex. Medial PFC representations related to item-level reinstatement in both groups, but this signature did not show developmental differences. Interestingly however, we did observe significant age differences in the organizational schemes predicting category-level reinstatement: Specifically, trial-by-trial hippocampal differentiation among children (but not adults) and mPFC integration among adults (but not children) each tracked with category reinstatement in VTC. These associations were unique to hippocampus and mPFC, as they were not observed in other regions showing significant integration in either children or adults (i.e., parietal cortex, posterior cingulate, and caudate). By testing the relationship between representations in hippocampus and mPFC and neocortical reinstatement, the current study further establishes a role for hippocampus and mPFC in reinstating neocortical traces in development—a correspondence that has previously only been shown in adults (Hindy, Ng & Turk-Browne, 2016; Mack & Preston, 2016). Collectively, our results reveal that children and adults both engage in neocortical memory reinstatement, but through different representational pathways, underpinned by hippocampal and mPFC organization.

Taken together, the present research provides key insight into the development of neural representations associated with learning, representing, and retrieving overlapping events from memory. We show that memory for related events was associated with different underlying representational schemes at different ages.

Specifically, children exhibited differentiation in the hippocampal system, which was linked to higher-fidelity neocortical reinstatement of event details, that in turn was associated with superior memory behavior. In contrast to children, hippocampal differentiation was unrelated to neocortical reinstatement in adults; instead, in adults mPFC integration benefitted neocortical reinstatement, which in turn tracked faster response speeds. Our findings align well with current perspectives of neural development which suggest that differences in the maturational pace of the hippocampus and prefrontal cortex—and the neural representations they support— contributes to the shift in balance between memory specificity and flexibility (Keresztes et al., 2018). Notably, the findings further suggest that children and adults form and retrieve memories using different types of representations. Importantly, given that the ability to organize, retrieve, and extend memories for related information is critical to knowledge development and predicts long-term academic success in both children and adults (Varga et al., 2019), the present results suggest that reducing overlap among related information may aid learning and memory in educationally relevant ways.

## Acknowledgments

The data reported here were collected with support by funding from NIMH R01 grant 100121 (ARP), NICHD F32 grant 095586 (NLV), NSERC Discovery Grant RGPIN-04933-2018 (MLS), Canada Foundation for Innovation JELF (MLS), and Ontario Research Fund 36876 (MLS). The funders had no role in study design, data collection and analysis, or preparation of the manuscript. The authors extend their gratitude to Katharine Guarino, Christine Coughlin, Kim Nguyen, Lauren Quesada, and Drew (Francis) Hussey for assistance with piloting, data collection and analysis. We thank Tammy Tran for assistance with stimulus development. We also thank Katherine Sherrill and Neal Morton for input on data processing. Finally, we thank the families and participants, as this work would not have been possible without their help.

## Author Contributions

M.L.S, M.L.M. and A.R.P. conceptualized the study. M.L.S., M.L.M., A.R.P., N.L.V. and H.E.R. designed the experimental task. N.L.V., H.E.R., L.M., and E.M.H. collected the data. N.L.V., M.L.S., M.L.M., and R.J.M. analyzed the data. N.L.V. drafted the paper, and M.L.S. and A.R.P. revised the manuscript for review and finalization by all other authors.

## Data Availability

The source data that support the findings of this study are provided in the Open Science Framework repository at https://osf.io/6ny4g/?view_only=8dc01c7fa3a6441b8267e9340024f712.

## Code Availability

The custom code—including all required content and instructions for use—that supports the findings of this study are available in the Open Science Framework repository at https://osf.io/6ny4g/?view_only=8dc01c7fa3a6441b8267e9340024f712.

