## Supplementary Information for "Differentiation of related events in hippocampus supports memory reinstatement in development"

**Supplementary Data**

**Age-related learning differences.** Most participants (n=47 out of 52) had a total of two learning exposures (Figure 1C). Five children required more than two exposures, yielding statistically more exposures on average in the child compared to the adult group (Child: *M* total exposures = 2.29, *SD* = 0.72; Adult: *M* total exposures = 2.00, *SD* = 0.00; *t*(26) = 2.13, *p* = .04, d=-0.57, 95% CI=[-0.58,-0.01]). Of the five children who required more than two exposures, three (one 8-year-olds and two 9-year-olds) required three exposures, one (7-year-old) required four exposures, and one (7-year-old) required five exposures. Notably, two of the children who had three exposures reached the 90% learning criterion on the first exposure, but then dropped below criterion on the second exposure. To rule out the possibility that their initial success was due to guessing rather than learning, we had the participants complete a third exposure. With these additional repetitions, children achieved a similar level of final learning performance to adults (Children: *Accuracy [Proportion correct]* = 0.96, *SD* = 0.03; Adults: *Accuracy [Proportion correct]* = 0.99, *SD* = 0.02). Nevertheless, performance on the final learning run was still statistically higher in adults, *t*(45.05) = 4.24, *p* < .001, d=1.16, 95% CI=[0.02, 0.04].

In addition to comparing learning performance between age groups, we also tested for age-related performance differences within the child group alone. More specifically, we tested for differences between younger (7- & 8-year-olds; *N* = 14) and older (9- & 10-year-olds, *N* = 13) children, which resulted in nonsignificant differences in initial learning (i.e., Learning Exposure 1; Younger: *Accuracy* = 0.83, *SD* = .10; Older: *Accuracy* = .82, *SD* = .18; *t*(18.69)=.19, *p*=.85, d=0.08, 95% CI=[-0.11,0.13]), final learning (Younger: *Accuracy* = 0.96, *SD* = .03; Older: *Accuracy* = .97, *SD* = .03; *t*(25)=-.59, *p*=.56, d=-0.23, 95% CI=[-0.03,0.02]), and the number of total learning exposures completed (Younger: *M total exposures* = 2.43, *SD* = .94; Older: *M total exposures* = 2.15, *SD* = .38; *t*(17.32) = 1.01, *p* =.33, d=0.38, 95% CI=[-0.29,0.85]).

Notably, most adults did not need a second learning exposure, but were required to complete one to better equate the number of total learning exposures across age groups. More specifically, whereas 76% of adults (*N* = 19) reached the 90% accuracy criterion on the first learning exposure, only 37% of children (*N* = 10 which includes the two children who dropped below criterion on the second exposure; see above) reached the same criterion within a single exposure. Thus, by requiring a minimum of two learning exposures regardless of initial performance, we ensured that most child and adult participants (n=47; see above) had an equivalent number of exposures, which would not have been the case otherwise.

**Hippocampal differentiation controlling for univariate activation.** The overall difference in pattern similarity between related and unrelated object cues observed in children in bilateral anatomical hippocampus (Figure 2b) remained when mean hippocampal univariate activation during the cue period was entered as a covariate into the analysis of variance, *F*(1,25)=4.65, *p*=.04, *η_p_*^2^ =.16, 95% CI=[-0.020,-0.002]. This finding suggests that the decreased representational similarity observed for related relative unrelated memories cannot be attributed to broad changes in mean hippocampal activation in response to the retrieval cue, but rather are the result of differences in the distributed patterns of activation across hippocampal voxels.

**Classifier validation accuracy.** To quantify the degree to which the target category was reinstated during the delay period, we used the category localizer to train a multivoxel pattern analysis (MVPA) classifier to identify patterns of activity associated with face and scene processing in the ventral temporal cortex (VTC) and parietal cortex functional regions of interest (fROIs; see Figure 3a). Ensuring that the MVPA classifier could accurately detect when participants were viewing faces and scenes in the category localizer task was a necessary first step before testing our key question, which was whether face and scene patterns were reinstated during the recall task (Figure 1b). Cross-validation performance was significantly above chance levels (Chance = 0.5; VTC: *M* = 0.97, *SD* = 0.02, *t*(51)=136.77, *p*<.001, d=18.97, 95% CI=[0.46,0.48], all participants above chance: *p*s < 1.50x10^-15^; parietal: *M* = 0.85, *SD* = 0.08, *t*(51)=32.62, *p*<.001, d=4.52, 95% CI=[0.33,0.37], all participants significantly above chance: *p*s < 7.70x10^-05^) and did not differ as a function of age group (VTC: *t*(50)=1.36, *p*=.18, d=0.38, 95% CI=[-0.004,0.023]; parietal: *t*(50)=1.36, *p*=.18, d=0.38, 95% CI=[-0.014,0.072]). Together, this pattern of results suggests that face and scene patterns were discriminable for each participant and age group in each fROI.

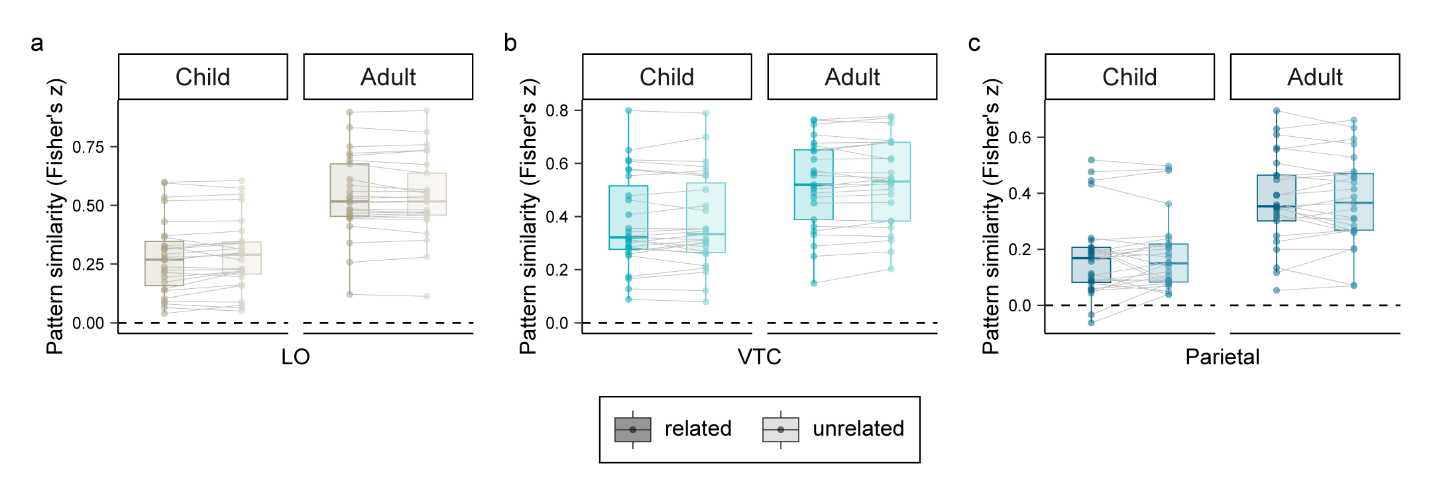

Supplementary Figure 1**. Children and adults show no evidence of neural organization (neither differentiation nor integration) in visual or reinstatement regions.** For **a-c**, plots feature pairwise similarity values averaged across object cues that were related or unrelated, separately for participants within each age group (n=52; 27 children, 25 adults). Box plots depict the median (middle line), 25^th^ and 75^th^ percentiles (box), and the largest individual values no greater than the 5^th^ and 95^th^ percentiles (whiskers). Dots reflect individual participant means, with dots extending beyond the whiskers reflecting outliers, defined as values that were 1.5 times greater than the interquartile range (IQR). **a** Neural patterns extracted from bilateral lateral occipital (LO) cortex, derived from an automated anatomical mask using Freesurfer segmentation. There was neither a main effect of similarity (*F*(1,50)=0.60, *p=*0.44*, η_p_*^2^=.01) nor an interaction with age group (*F*(1,50)=2.43, *p=*0.13*, η_p_*^2^=.05). Given that significant hippocampal differentiation was only observed in the child group, we also examined the effect of similarity in this group alone. There were no significant differences between related and unrelated object cues (*F*(1,26)=1.87, *p=*0.18*, η_p_*^2^=.07, 95% CI=[-0.02,0.01]), suggesting that the primary hippocampal differentiation effects cannot be attributed to neural similarity of the visual objects themselves, but rather result from mnemonic coding schemes. **b** Neural patterns extracted from functionally-defined ventral temporal cortex (VTC; see Figure 3a and Methods). There was neither a main effect of similarity (*F*(1,50)=0.88, *p=*0.35*, η_p_*^2^=.02) nor an interaction with age group (*F*(1,50)=0.85, *p=*0.36*, η_p_*^2^=.02). There were also no significant difference between related and unrelated object cues when we examined children alone, (*F*(1,26)<0.001, *p=*0.99*, η_p_*^2^<.001, 95% CI=[-0.015,0.014]). **c** Neural patterns extracted from functionally-defined parietal cortex (Figure 3a and Methods). There was neither a main effect of similarity (*F*(1,50)=0.28, *p=* 0.59*,* *η_p_*^2^=.01) nor an interaction with age group (*F*(1,50)=0.52, *p=*0.48*, η_p_*^2^=.01). There were also no significant difference between related and unrelated object cues when we examined children alone (*F*(1,26)=0.59, *p=*0.45*, η_p_*^2^=.02, 95% CI=[-0.041,0.019]).

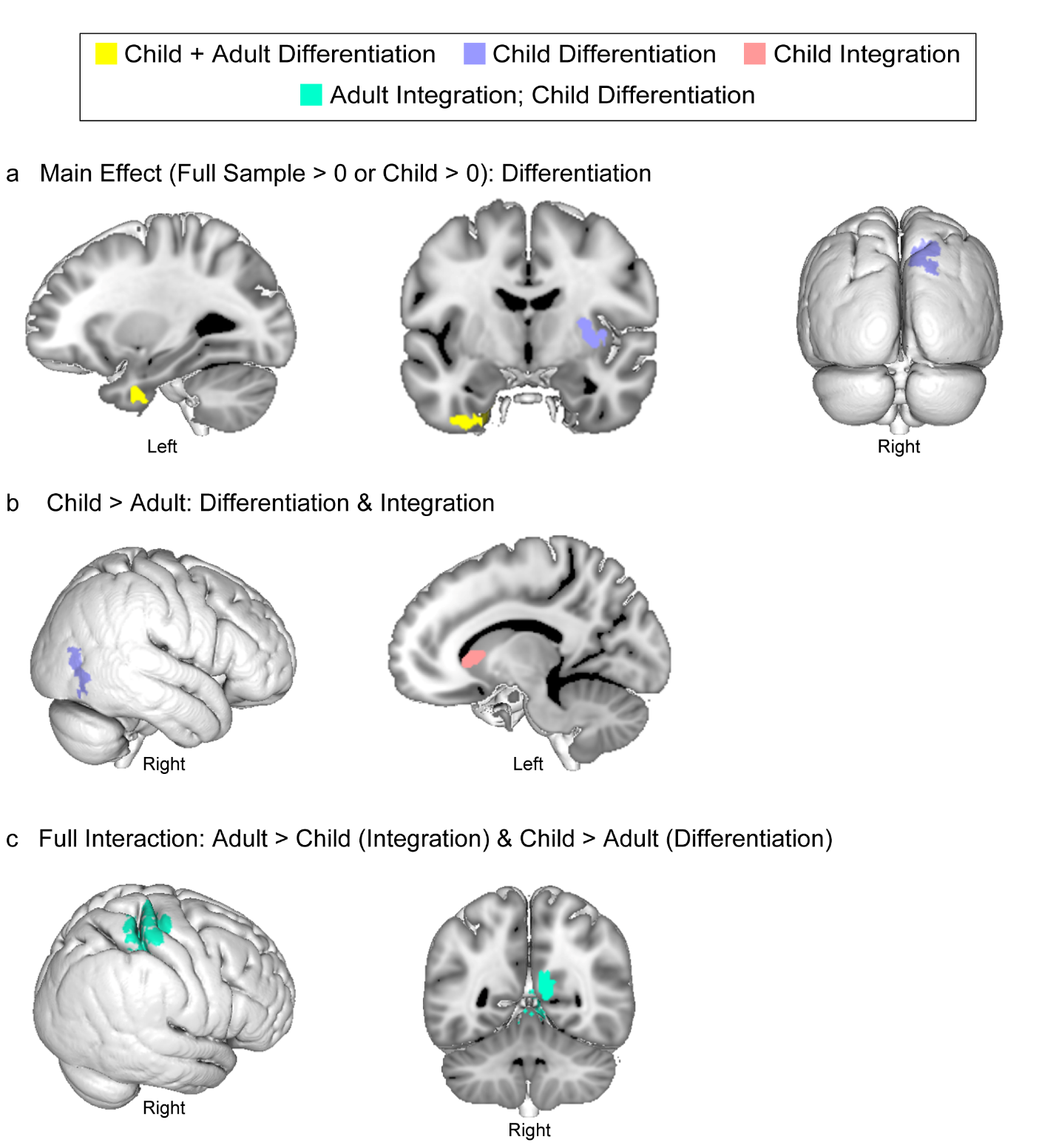
Supplementary Figure 2. **Whole-brain RSA cue integration/differentiation searchlight results.** **a** Regions showing a main effect of differentiation, either across the full sample (yellow) or only in children (purple) when the cluster identified in the full sample was interrogated in each age group separately (as in Figure 2c-d). **b** Regions showing developmental differences in differentiation (purple) and integration (pink), which were evidenced in children but not adults (Child > Adult). **c** Regions showing an interaction between coding scheme and age group (teal), such that adults showed significant integration but children show significant differentiation in the same voxels simultaneously. For **a-c**, clusters displayed on the 1-mm MNI template brain. See also Supplementary Figure 3 and Supplementary Table 2.

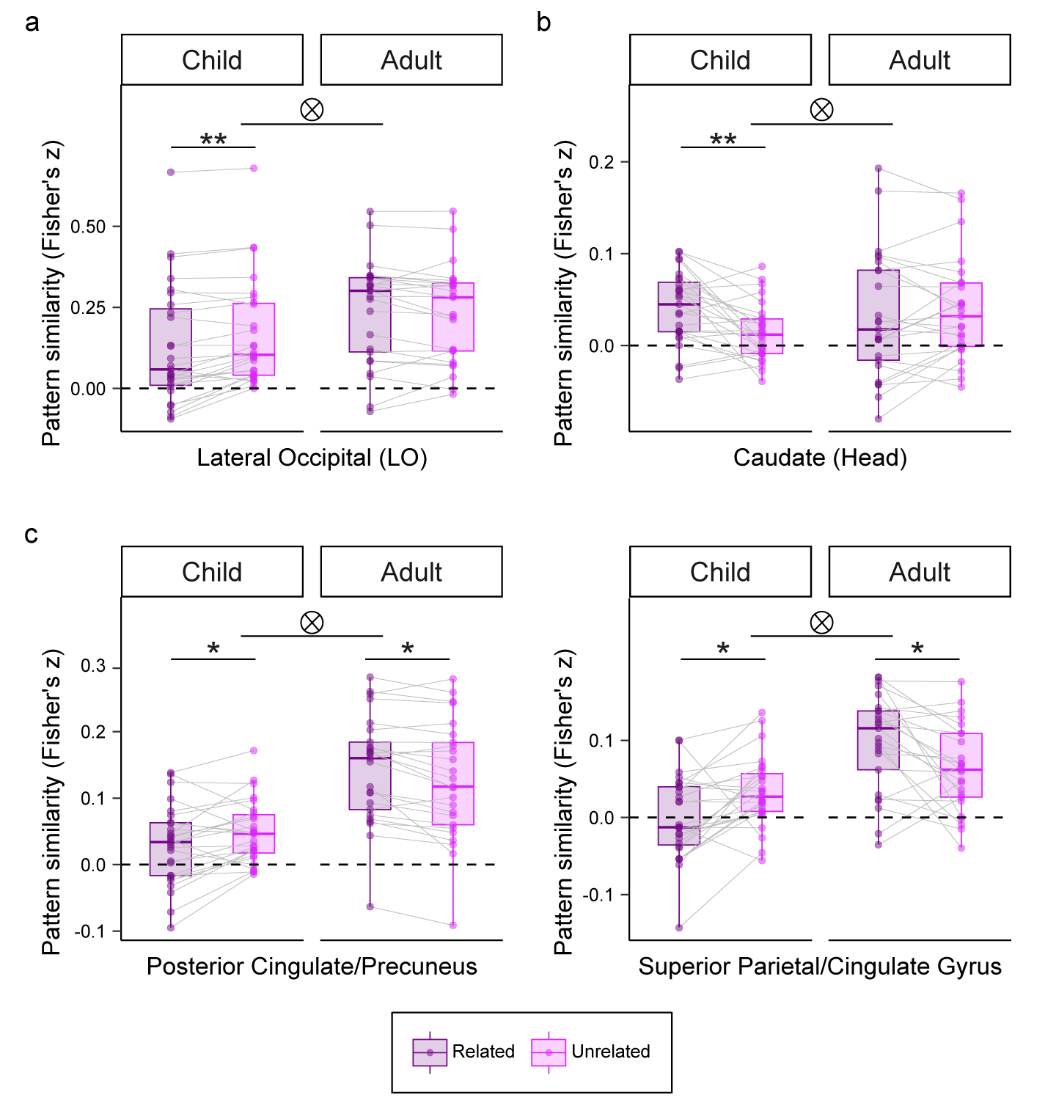

Supplementary Figure 3. **Neural patterns extracted from whole-brain RSA searchlight regions that showed significant developmental differences.** We compared the similarity of fMRI patterns for related as compared with unrelated objects across trials in regions for which there was an age-related difference in differentiation (related < unrelated) or integration (related > unrelated) (from Supplementary Figure 2 & Supplementary Table 2). Boxplots depict regions for which, when we tested effects within each age group separately, we found significant **a** differentiation or **b** integration within at least one group (here, children), or **c** significant but opposite coding schemes between age groups (here, integration in adults but differentiation in children). For **a-c**, plots feature pairwise similarity values averaged across object cues that were related or unrelated, separately for participants within each age group (n=52; 27 children, 25 adults). Box plots depict the median (middle line), 25^th^ and 75^th^ percentiles (box), and the largest individual values no greater than the 5^th^ and 95^th^ percentiles (whiskers). Dots reflect individual participant means, with dots extending beyond the whiskers reflecting outliers, defined as values that were 1.5 times greater than the interquartile range (IQR). Asterisks reflect a significant related/unrelated difference within an age group at a threshold of * p < .05 or ** p < .001. Tensor product symbols indicate a significant interaction between age group and related/unrelated conditions (marked here simply for clarity).

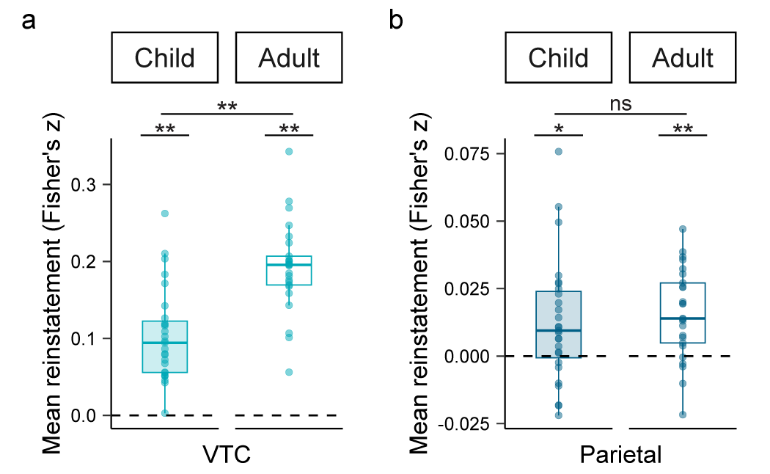

Supplementary Figure 4. **Children and adults show category reinstatement using an RSA approach.** Representational similarity analysis was implemented at the level of the full category-selective functional regions of interest (fROI; see Figure 3a). For each scanned recall trial, we correlated fMRI activation patterns from the 9-second delay period with viewing of the same category during a pre-exposure item perception localizer (same-category), which was compared to viewing of items from a different visual category (different-category baseline). For each participant, we computed a single category reinstatement index by averaging the mean same category – different category similarity difference across recall trials. **a** VTC and **b** parietal cortex category reinstatement. A reinstatement index reliably above 0 (i.e., the black dashed line) indicates significant target category evidence. Box plots depict the median (middle line), 25^th^ and 75^th^ percentiles (box), and the largest individual values no greater than the 5^th^ and 95^th^ percentiles (whiskers). Dots reflect individual participant means; n = 52 per plot (27 children; 25 adults), with dots extending beyond the whiskers reflecting outliers, defined as values that were 1.5 times greater than the interquartile range (IQR). Asterisks above each box reflect a significant within-group effect at a one-tailed threshold of p < .05 (*) or p < .001 (**), while asterisks between bars reflect a significant between-group effect at a two-tailed threshold of **p < .001 or that are non-significant (ns).

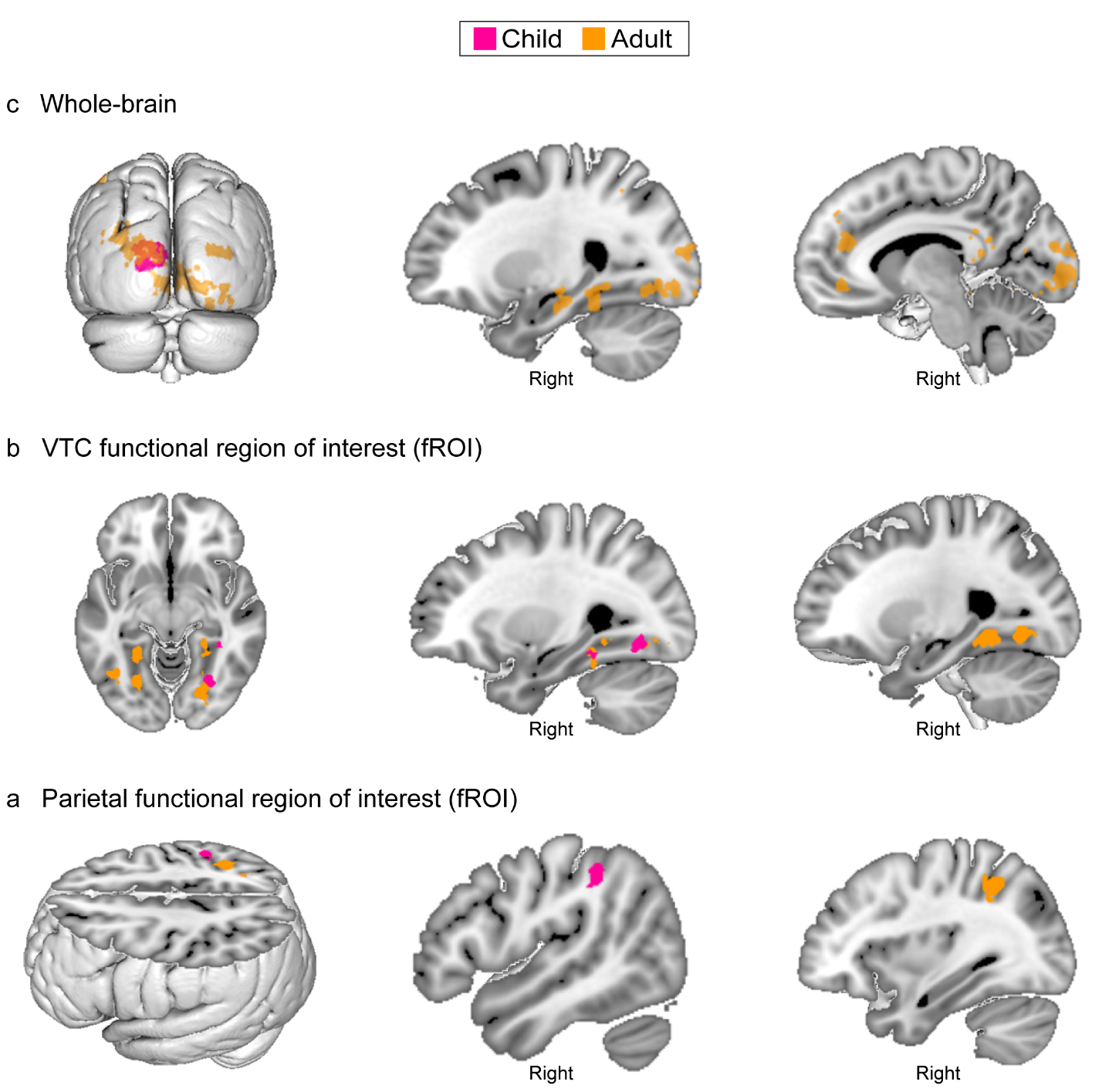

Supplementary Figure 5. **Whole-brain RSA item reinstatement searchlight results.** To complement the main functional regions of interest (fROI) item reinstatement analyses (Figure 4b), we searched for voxels that exhibited significant item reinstatement at the whole brain level, testing for this effect within in the child (pink) and adult (orange) groups separately (i.e., Child > 0; Adult > 0). **a** Regions showing significant item reinstatement at the whole-brain level. **b-c** Regions showing significant item reinstatement when they were small-volume corrected within the fROIs in **b** VTC and **c** parietal cortex. For **a-c**, clusters displayed on the 1-mm MNI template brain. See also Supplementary Table 3 and Supplementary Figure 6 for corresponding results in hippocampus.

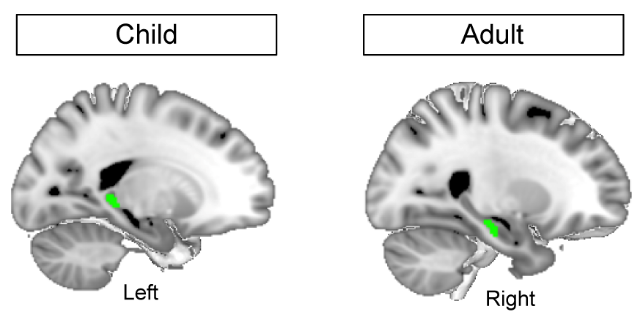

Supplementary Figure 6. **Children and adults reinstate target items in different subregions of hippocampus.** Results of item reinstatement representational similarity analysis approach (Figure 4a; see also Supplementary Table 3 and Supplementary Figure 5), which was implemented at the individual voxel level via separate searchlight analyses in children (child > 0) and adults (adult > 0). For each participant and voxel, we computed a single item reinstatement index by averaging the mean same item – different item similarity difference across recall trials. We observed that children reinstated specific memory associates in posterior hippocampus (*x*, *y*, *z* = -18.7, -37.5, -0.67), whereas adults exhibited specific item representations in anterior hippocampus (*x*, *y*, *z* = 28.3, -20.7, -17.2; Fig. 4c). This mirrors the general pattern that has been observed previously in terms of locus of univariate activation shifting from posterior to anterior hippocampus across development during associative retrieval^34^. These results thus converge with the view that memory processes are differently distributed in the immature hippocampus^50^. By identifying the specific parts of hippocampus associated with organization of related memories (Figure 2b & 2c) from those involved in reinstatement of specific memory features here, we build on this proposal, showing that the early hippocampus prioritizes memory specificity, through disambiguating similar events via differentiation while simultaneously retrieving item-specific traces in posterior regions. Significant item reinstatement clusters are displayed on the 1-mm MNI template brain. Clusters were identified on a whole-brain template (see Supplementary Table 3 and Supplementary Figure 5 for additional regions) and are significant after small-volume correction for multiple comparisons within anatomical hippocampus.

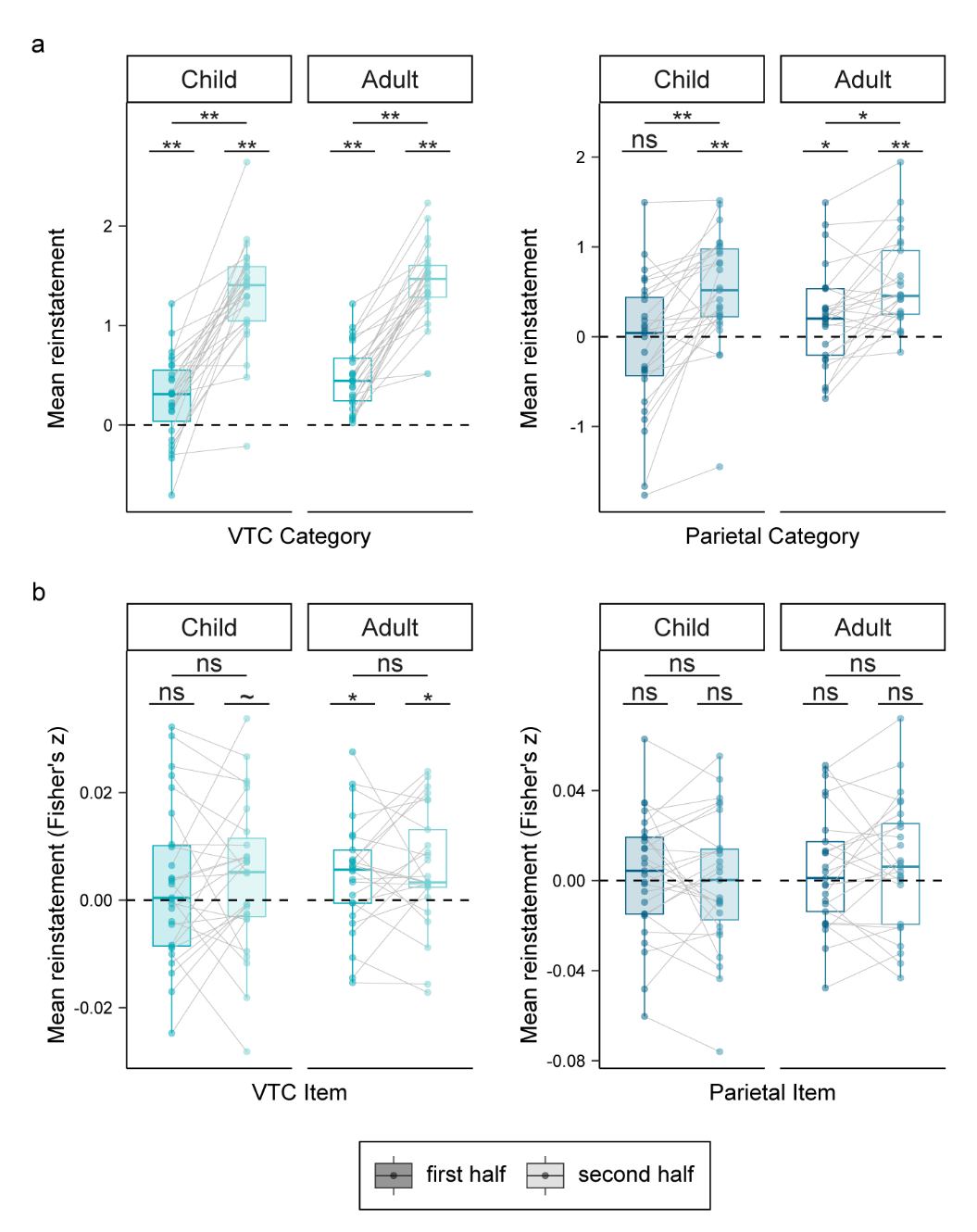

Supplementary Figure 7. **Reinstatement generally increased across the delay in children and adults.** **a** At the category level, reinstatement in VTC and parietal cortex significantly increased from the first to second half of the delay for children (VTC: *p*<.001; parietal: *p*<.001) and adults (VTC: *p*<.001; parietal: *p*=.001). Moreover, category-level reinstatement was significant (i.e., above 0) in both the first and second halves of the delay in VTC for both children and adults (*p*s≤.001) and in parietal for adults (*p*s<.03). In contrast, for children, parietal reinstatement was significant in the second half (*p*<.001) but not the first half (*p*=.26) of the delay. Moreover, age group differences were observed in the first half of the delay (at trend or significant levels; VTC: *t*(50)=2.04, *p*=.047, d=0.57, 95% CI=[0.003,0.428]; parietal: *t*(50)=1.71, *p*=.09, d=0.47, 95% CI=[-0.06,0.69]) but not during the second half of the delay (VTC: *t*(50)=.77, *p*=.44, d=0.22, 95% CI=[-0.16,0.36]; parietal: *t*(50)=.39, *p*=.70, d=0.11, 95% CI=[-0.26,0.38]). These results suggest that reinstatement increased across the delay, with children being slower to reinstate memories. By the end of the delay, age differences were largely eliminated. **b** Item reinstatement did not significantly differ between the first and second halves of the delay in either children or adults (*p*s>.51). Moreover, children and adults did not differ from one another in the second half (all |*ts*(50)|<0.89, all *ps*>.37), nor in the first half (all |*ts*(50)|<0.54, all *p*s>.59) of the delay. However, considering reinstatement within each group separately by half revealed that while adults showed significant item reinstatement in both the first (*p*=.02) and second (*p*=.006) halves of the delay in VTC, children showed a trend for this effect only during the second half (*p* = .06; not the first: *p* = .16). Item reinstatement was not significant in either delay interval or at either age in parietal cortex (*t*s<1.19; *p*s>.12). For **a**-**b**, Asterisks above each box reflect a significant within-group effect at a one-tailed threshold of p<.05(*) or p≤.001(**), while asterisks between boxes reflect a significant between-group effect at a two-tailed threshold of p<.05(*), p<.001(**), or that are at trend (~) or non-significant (ns) levels.

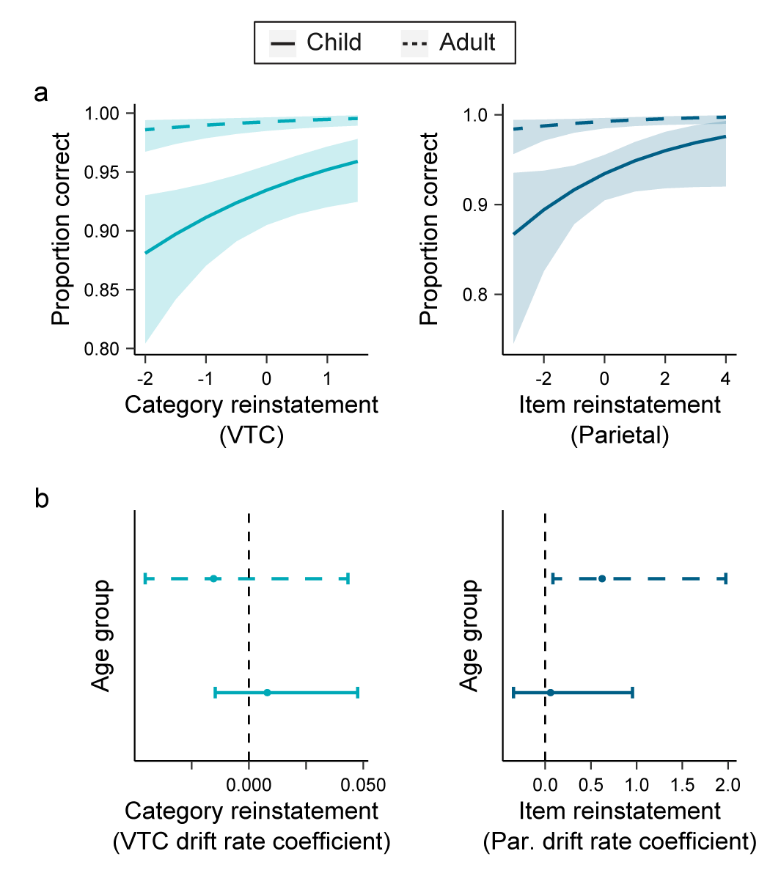

Supplementary Figure 8. **Reinstatement during the second half of the delay was behaviorally significant in children and adults.** Having found that generally reinstatement was stronger toward the end of the delay (Supplementary Figure 7), we were additionally curious as to whether this end-of-delay reinstatement would be related to behavior. Re-running our mixed-effects analyses (Figure 5a) and drift-diffusion models (DDM; Figure 5b) separately considering reinstatement evidence restricted to the second half of the delay period replicated our main brain-behavior results. Specifically, **a** more evidence for category reinstatement in VTC (χ^2^_1_=5.65, *p*=.03) and a trend for item reinstatement in parietal cortex (χ^2^_1_=3.58, *p*=.06) during the second half of the delay was related to more accurate responses for children; the same was not true for the first-half reactivation values (*p*s>0.17; not depicted). Here, significant predictors are displayed in separate plots for visualization purposes only, as all predictors (see Method) were included in a single statistical model. All predictors were scaled and centered within participants to remove participant-specific effects. For both plots, n = 52 (27 children; 25 adults); age group was modeled as a main effect. **b** In addition, evidence of item reinstatement in parietal cortex during the second half of the delay period was related to response speed on match trials in adults (*p*=.03); the same was not true for the first-half reinstatement values (*p*s > .19). Notably, we did not find a significant association for item reinstatement in parietal or category reinstatement in VTC with response speed during either half of the delay interval in children (*p*s > 0.28; first half of delay not depicted). Here, the VTC category and parietal cortex item reinstatement indices were entered into a four separate DDM models of response probability and speed, one each for each delay half (first versus second) which was further separated by children and adults. The lines represent 95% confidence intervals of the 2,000 posterior parameter estimates (one-tailed), with circles representing the mean of the posterior parameter distributions. Statistical significance was determined by calculating the proportion of samples that were less than 0, equivalent to a one-tailed test.

**Supplementary Table 1.** Mean and range of number of correct trials for final learning, scanned recall, and for the overall neural analyses after accounting for learning and scanned recall, separately for each age group.

|  | Learning/Retrieval | |  | Scanned Recall | |  | Overall | |
| --- | --- | --- | --- | --- | --- | --- | --- | --- |
|  | *Mean (SD)* | *Range* |  | *Mean (SD)* | *Range* |  | *Mean (SD)* | *Range* |
| Child | 34.67 (1.07) | 33-36 |  | 30.96 (3.26) | 25-35 |  | 29.96 (2.90) | 25-35 |
| Adult | 35.68 (0.69) | 33-36 |  | 35.32 (0.95) | 33-36 |  | 35.08 (1.32) | 31-36 |

*Note: The minimum number of trials represented in the table (25) corresponds with the behavioral threshold for above-chance (0.5) performance on scanned recall, based on a Binomial Test (p<.05; two-tailed).*

**Supplementary Table 2.** Whole-brain RSA cue differentiation/integration searchlight results. Regions reflect clusters of voxels that survived cluster correction. For clusters showing a significant age group difference in coding scheme, effects were considered significant if cue differentiation/integration was reliable at the within-group level in the group that showed the enhanced pattern. Values (X, Y, Z) reflect cluster center of gravity (COG).

|  | *Region* | *Hemisphere* | *N vox* | *t* | *X* | *Y* | *Z* |
| --- | --- | --- | --- | --- | --- | --- | --- |
| **Differentiation** | ***Main Effect*** | | | | | | |
|  | Cuneus^a^ | R | 164 | 1.84 | 13 | -81 | 39 |
|  | Anterior parahippocampal gyrus^e^ | L | 120 | 2.37 | -29 | -5 | -41 |
|  | Insula cortex^a^ | R | 118 | 3.51 | 34 | -7 | 3 |
|  | Subgenual anterior cingulate cortex^a^ | L | 99 | 3.54 | -6 | 6 | -9 |
|  | ***Adult > Child*** | | | | | | |
|  | None | - | - | - | - | - | - |
|  | ***Child > Adult*** | | | | | | |
|  | Posterior cingulate/precuneus^c^ | R | 568 | 2.91 | 8 | -43 | 6 |
|  | Superior parietal lobule/cingulate gyrus^c^ | R | 482 | 1.54 | 31 | -34 | 57 |
|  | Lateral occipital cortex^a^ | R | 124 | 2.77 | 56 | -63 | 1 |
| **Integration** | ***Main Effect*** | | | | | | |
|  | None | - | - | - | - | - | - |
|  | ***Adult > Child*** | | | | | | |
|  | Posterior cingulate/precuneus^d^ | R | 568 | 2.91 | 8 | -43 | 6 |
|  | Superior parietal lobule/cingulate gyrus^d^ | R | 482 | 1.54 | 31 | -34 | 57 |
|  | Medial prefrontal cortex (mPFC)^b^ | L | 188 | 4.14 | -15 | 45 | -4 |
|  | ***Child > Adult*** | | | | | | |
|  | Caudate (head)^c^ | L | 142 | 3.61 | -11 | 21 | 3 |

*Note: 2 mm isotropic voxels. MNI coordinates rounded to the nearest mm. Age difference patterns within clusters that showed a main effect or age group difference: ^a^differentiation in children, null pattern in adults, ^b^integration in adults, null pattern in children, ^c^integration in children, null pattern in adults, ^d^integration in adults, differentiation in children, ^e^trend for differentiation in both groups (ps≤.07). See also Supplementary Figures 2 & 3 for corresponding whole-brain visualizations and within-group plots, respectively.*

**Supplementary Table 3.** Whole-brain RSA item reinstatement searchlight results. Regions reflect clusters of voxels that survived cluster correction within each age group separately. Values (X, Y, Z) reflect cluster center of gravity (COG).

| *Region* | *Hemisphere* | *N vox* | *t* | *X* | *Y* | *Z* |
| --- | --- | --- | --- | --- | --- | --- |
| ***Child*** | | | | | | |
| Occipital pole | L | 112 | 1.71 | -11 | -100 | 11 |
| Intraparietal sulcus/inferior parietal lobule^┼^ | R | 61 | 3.29 | 48 | -38 | 44 |
| Fusiform* | R | 39 | 3.28 | 28 | -67 | -9 |
| Parahippocampal cortex* | R | 23 | 1.99 | 34 | -39 | -12 |
| ***Adult*** | | | | | | |
| Occipital lobe** | B | 1064 | 2.86 | 2 | -92 | 4 |
| Dorsomedial prefrontal ctx/ant. cingulate | B | 370 | 2.58 | -2 | 42 | 24 |
| Posterior cingulate/medial occipital | B | 275 | 2.36 | -1 | -55 | 17 |
| Medial prefrontal cortex (mPFC) | B | 192 | 2.4 | 2 | 53 | -5 |
| Intraparietal sulcus^┼┼^ | R | 135 | 2.96 | 32 | -51 | 44 |
| Anterior hippocampus/MTL cortex | R | 111 | 1.82 | 34 | -23 | -20 |
| Angular gyrus | L | 109 | 2.53 | -49 | -51 | 20 |
| Posterior parahippocampal/fusiform gyrus** | R | 101 | 2.07 | 26 | -41 | -12 |
| Fusiform gyrus** | L | 93 | 5.28 | -23 | -68 | -8 |
| Inferior frontal gyrus/Insula | R | 91 | 2.15 | 38 | 24 | -5 |
| Lingual/fusiform gyrus** | L | 89 | 2.57 | -23 | -48 | -10 |
| Superior parietal lobule | L | 88 | 3.85 | -47 | -37 | 57 |
| Frontal pole | L | 82 | 2.09 | -27 | 54 | 11 |
| Lateral occipital* | L | 63 | 2.95 | -40 | -63 | -6 |
| Lateral occipital* | R | 49 | 3.98 | 40 | -81 | -16 |

*Note: 2 mm isotropic voxels. MNI coordinates rounded to the nearest mm. *denotes significant after small volume correction in VTC; **denotes significant in whole-brain and VTC small-volume corrected analyses;* ^┼^*denotes significant after small volume correction in parietal cortex;* ^┼┼^*denotes significant in whole-brain and parietal small-volume corrected analyses. See also Supplementary Figures 5 & 6 for corresponding whole-brain and hippocampal cluster visualizations, respectively.*
